# Soma to neuron communication links stress adaptation to stress avoidance behavior

**DOI:** 10.1101/2025.05.07.652728

**Authors:** Julia Witrado, Ella Gundrum, Maria Victoria Veroli, A.J. Gilsrud, Mahdi Eskandarisani, Ioannis K. Zervantonakis, Steven Mullett, Stacy Gelhaus, Todd Lamitina

## Abstract

In multicellular organisms, signaling from the nervous system to the peripheral tissues can activate physiological responses to stress. Here, we show that inter-tissue stress communication can also function in reverse, i.e. from the peripheral tissue to the nervous system. *osm-8* mutants, which activate the physiological osmotic stress response in the *C. elegans* skin, also exhibit defective osmotic avoidance (Osm) behavior, via a direct and specific effect on ASH osmosensory neuron excitability. Both *osm-8* and the Patched-related gene *ptr-23*, mutations in which suppress all *osm-8* phenotypes, function in the hypodermal lysosomes to regulate both physiology and behavior. Unbiased lipidomics shows that *osm-8* leads to a *ptr-23*-dependent elevation of the lysosome specific lipid bis(monoacylglycero)phosphate (BMP) and expansion of the pool of hypodermal lysosomes. Just as genetic activation of the osmotic stress response by loss of *osm-8* in the hypodermis causes an Osm phenotype, acute physiological exposure to osmotic stress also confers a reversible Osm phenotype. Behavioral and genetic plasticity requires biosynthesis of the compatible solute glycerol, a key physiological output of the organismal osmotic stress response. However, *ptr-23* is only required for *osm-8* induced behavioral plasticity and not physiological plasticity. Instead, both genetic and physiologically induced Osm phenotypes require the unusual non-neuronal lysosomal V-ATPase subunit *vha-5*, which is also critical for organismal osmotic stress survival. Together, these data reveal that genetic or physiological activation of stress signaling from the skin elicits lysosome-associated signals that modulate organismal neurophysiology to attenuate a sensory neuron circuit. Such ‘body-brain’ interoceptive communication may allow organisms to better match neuronal decision-making with organismal physiological state.

## Introduction

### Sensory neuron control of somatic stress responses

Cellular responses to alterations in homeostasis, such as changes in temperature, oxygen, and osmolarity, are commonly referred to as cellular stress responses. This is because cells contain autonomous mechanisms for sensing, signaling, and adapting to the particular challenges posed by each individual stressor. However, in an organismal setting, these autonomous cellular stress response pathways become subservient to systemic control mechanisms. For example, in mice, genetic or pharmacological activation of autophagy in the hypothalamic POMC neurons leads to activation of autophagy in peripheral tissues such as the liver and adipose cells (Martinez-Lopez *et al*. 2016). In *C. elegans*, genetic activation of the mitochondrial or endoplasmic reticulum stress response in neurons leads to activation of these same stress response pathways in peripheral intestinal cells (Taylor and Dillin 2013; Berendzen *et al*. 2016). Similarly, optogenetic activation of temperature-sensing neurons can activate the heat shock transcription factor in the germline (Tatum *et al*. 2015). While this neuron-to-soma control of stress response pathways is now well established, what is not clear is whether or not information flows in the reverse direction, i.e. from soma to neurons. Some studies suggest that food and/or pathogen based stress signaling from epithelial cells in *C. elegans* can signal to neurons to modify physiological states, such as longevity, sleep, and learned pathogen avoidance (Sinner *et al*. 2021; Savini *et al*. 2022; Wu *et al*. 2025). Whether or not abiotic factors, such as osmolarity, also modify behavior through changes in physiological state is less well understood.

### Osmotic behavior and physiology in *C. elegans*

The nematode *C. elegans* exhibits two robust responses to increased environmental osmolarity, i.e. hypertonic stress. First, *C. elegans* exhibit behavioral avoidance to hypertonicity (Culotti and Russell 1978). This response is driven by a well-described neuronal avoidance circuit made up of the amphid and phasmid sensory neurons ASH and PHB which signal to the command interneuron AVA to initiate backwards motion and reorientation away from the stressor (Kaplan and Horvitz 1993; Gat *et al*. 2023). Second, when worms are unable to avoid hypertonicity, they activate physiological adaptation pathways, primarily in the hypodermis (Lamitina *et al*. 2004). This includes transcriptional up- and down-regulation of >300 osmotic response genes, such as the glycerol biosynthetic enzyme *gpdh-1* (Rohlfing *et al*. 2010) via the activity of hypodermal molecular signaling pathways (Urso *et al*. 2020; Urso *et al*. 2023). Glycerol accumulation closely tracks extracellular solute levels which allows *C. elegans* to maintain cell and tissue volume without raising cytoplasmic ionic strength (Lamitina *et al*. 2004). Currently, the connection between osmotic avoidance behavior and peripheral tissue osmotic stress responses are not clear. ASH osmo-sensing does not appear to be required for hypertonicity induced *gpdh-1* upregulation or glycerol accumulation in the somatic tissues (Lee *et al*. 2016). Whether or not somatic stress signaling modifies ASH osmotic avoidance behavior has not been investigated.

### osm-8 and ptr-23

Upregulation of the glycerol biosynthetic enzyme *gpdh-*1 in the epithelial tissues is a robust and specific marker of the physiological response to hypertonic stress in *C. elegans* (Lamitina *et al*. 2006). *gpdh-1* upregulation by hypertonicity is positively regulated by the O-GlcNAc transferase *ogt-1* and the polyadenylation and cleavage factor complex subunits such as Symplekin/*symk-1* (Urso *et al*. 2020; Urso *et al*. 2023). Unexpectedly, *gpdh-1* is also under extensive negative regulation, as inhibition of >100 genes cause significant upregulation of *gpdh-1* under non-stressful isotonic conditions (Lamitina *et al*. 2006). These negative regulators result in a physiological state change, as mutant animals become completely resistant to dehydration when exposed to extreme hypertonicity (>500mM NaCl) and accumulate the osmolyte glycerol at close to molar levels (Solomon *et al*. 2004; Wheeler and Thomas 2006). One of these negative regulators is *osm-8*, a 331 residue protein with a signal sequence and S/T-rich repeats (Rohlfing *et al*. 2011). Loss of *osm-8* leads to upregulation of *gpdh-1*, as well as most of the ORG transcription program, and accumulation of glycerol. As a result, >90% of *osm-8* mutants retain mobility in >500mM NaCl, a condition in which 0% of wild type animals are motile. Through a genetic suppressor screen, we found that all of these physiological phenotypes require the Nieman-Pick C1 (NPC1)/patched-related (ptr) receptor *ptr-23* (Rohlfing et al. 2011). Tissue-specific and temporal rescue experiments reveal that *osm-8* is required in the hypodermal epithelial cells during the L4 stage for these phenotypes. If and how *osm-8* and *ptr-23* regulate neuronal osmotic avoidance behavior has not been determined.

Here, we show that, in addition to its role in regulating peripheral physiology, *osm-8* also regulates neuronal osmotic avoidance behavior by reducing the sensitivity of the ASH neuron to hypertonic stimuli. *osm-8* behavioral defects are completely dependent on *ptr-23*. Furthermore, the expression of *osm-8* and *ptr-23* in the hypodermis is necessary and sufficient to regulate neuronal behavior. Endogenously tagged alleles show that OSM-8 and PTR-23 co-localize to the hypodermal lysosomes. Unbiased lipidomic analysis shows that the lysosome-specific lipid bis(monoacylglycero)phosphate (BMP) exhibits *ptr-23*-dependent upregulation in *osm-8* mutants, which results in an expansion of the pool of lysosomes in the hypodermis. Acute adaptation of wild type worms to mild hypertonicity also induces a reversible osmotic avoidance defective phenotype like *osm-8*. The acquired Osm phenotype is dependent on both glycerol biosynthesis enzymes *gpdh-1* and *gpdh-2* and the lysosomal V-ATPase subunit *vha-5*, which is also required for *osm-8* behavioral phenotypes. Together, these data suggest that the *osm-8-ptr-23-vha-5* pathway that regulates peripheral stress physiology in the skin also modulates ASH-driven behavioral sensitivity to osmotic stress, thus synchronizing behavioral and physiological states at the organismal level.

## Results

### *osm-8* and *ptr-23* regulate osmotic stress resistance and osmotic behavioral avoidance

The single *osm-8* allele, *n1518*, was previously shown to constitutively upregulate the glycerol biosynthesis gene *gpdh-1*, contain elevated glycerol levels, and exhibit an osmotic stress resistance (Osr) phenotype to 500 mM NaCl, a normally lethal hypertonic stress to naïve wild type animals (Rohlfing *et al*. 2011). All of these phenotypes are dependent on the function of the patched-related receptor *ptr-23*. To create definitive null alleles for these genes, we generated deletion alleles for both *osm-8* and *ptr-23* using CRISPR/Cas9 (Fig. 1A). Similar to the originally reported alleles, the *osm-8(dr170)* allele was also Osr while the *ptr-23(dr180)* allele exhibited no Osr phenotype on its own but fully suppressed the Osr phenotype of *osm-8(d170)* (Fig. 1B). Mutations in neuron-expressed Osm genes, such as *osm-6*, do not exhibit an Osr phenotype (Fig. 1B). These new alleles represent definitive null alleles for both *osm-8* and *ptr-23* that phenocopy previously described mutants and were used for all subsequent studies, unless otherwise indicated.

**Figure 1.**
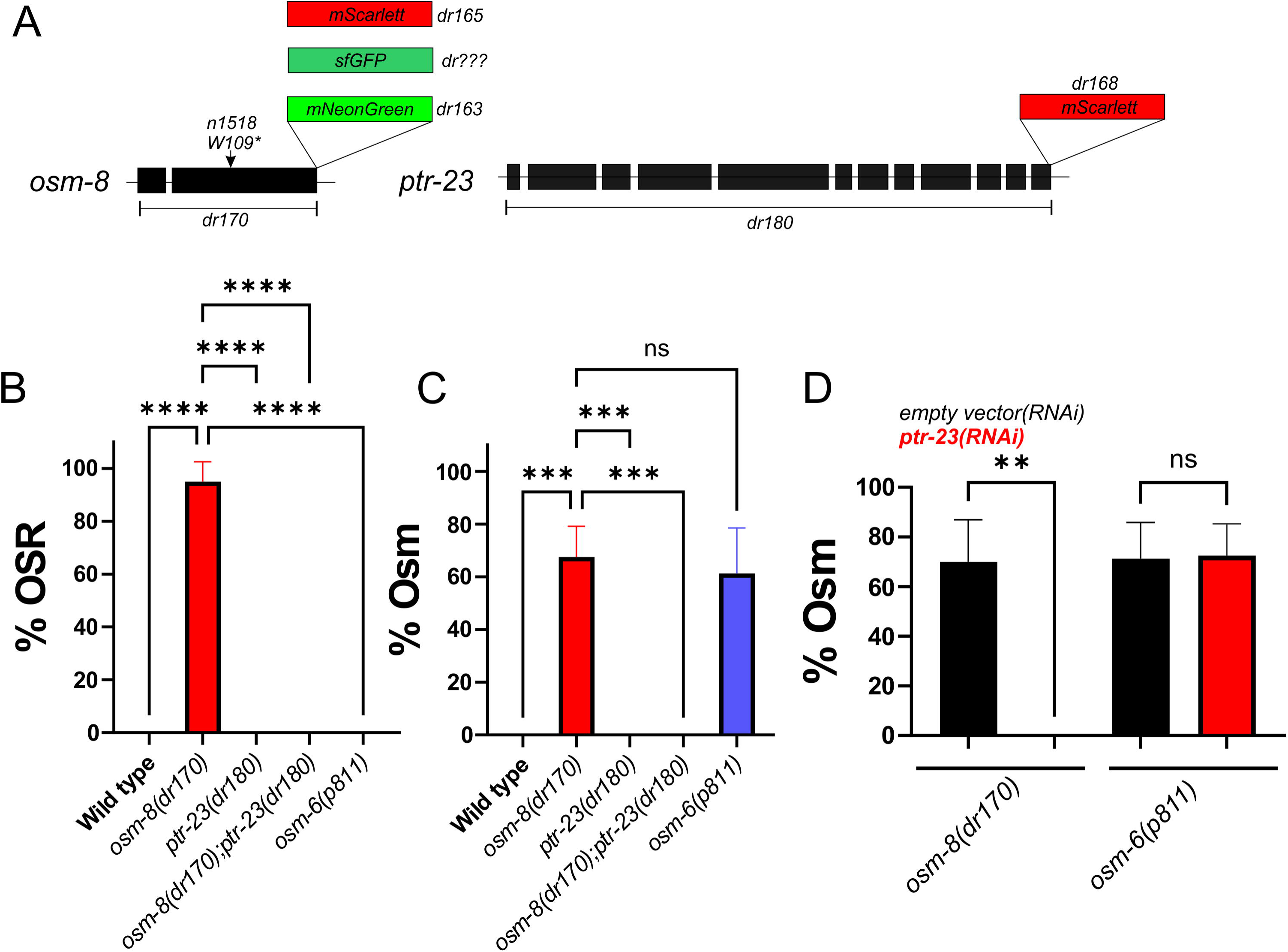
*osm-8* physiological and behavioral phenotypes depend on *ptr-23*. A) Diagram of the *osm-8* and *ptr-23* alleles generated in this study. A detailed list of these and other strains can be found in Table 1. The *osm-8(n1518)* allele was characterized in a prior study (Rohlfing *et al*. 2011). B) Osmotic stress resistance (OSR) phenotype of the indicated genotypes. N=8 replicates per genotype (10 animals per replicate, N=80 per genotype). ****-p<0.0001, One-way ANOVA with Kruskal-Wallis post hoc test. Individual data points are shown along with the mean ± S.D. C) Osmotic avoidance phenotype of the indicated genotypes. N=8 replicates per genotype (10 animals per replicate, N=80 per genotype). ***-p<0.001, ‘ns’ – p>0.05, One-way ANOVA with Kruskal-Wallis post hoc test. Individual data points are shown along with the mean ± S.D. D) Osmotic avoidance phenotype of the indicated genotypes. N=8 replicates per genotype (10 animals per replicate, N=80 per genotype). **-p<0.01, ‘ns’ – p>0.05, One-way ANOVA with Kruskal-Wallis post hoc test. Individual data points are shown along with the mean ± S.D.

**Table 1.**
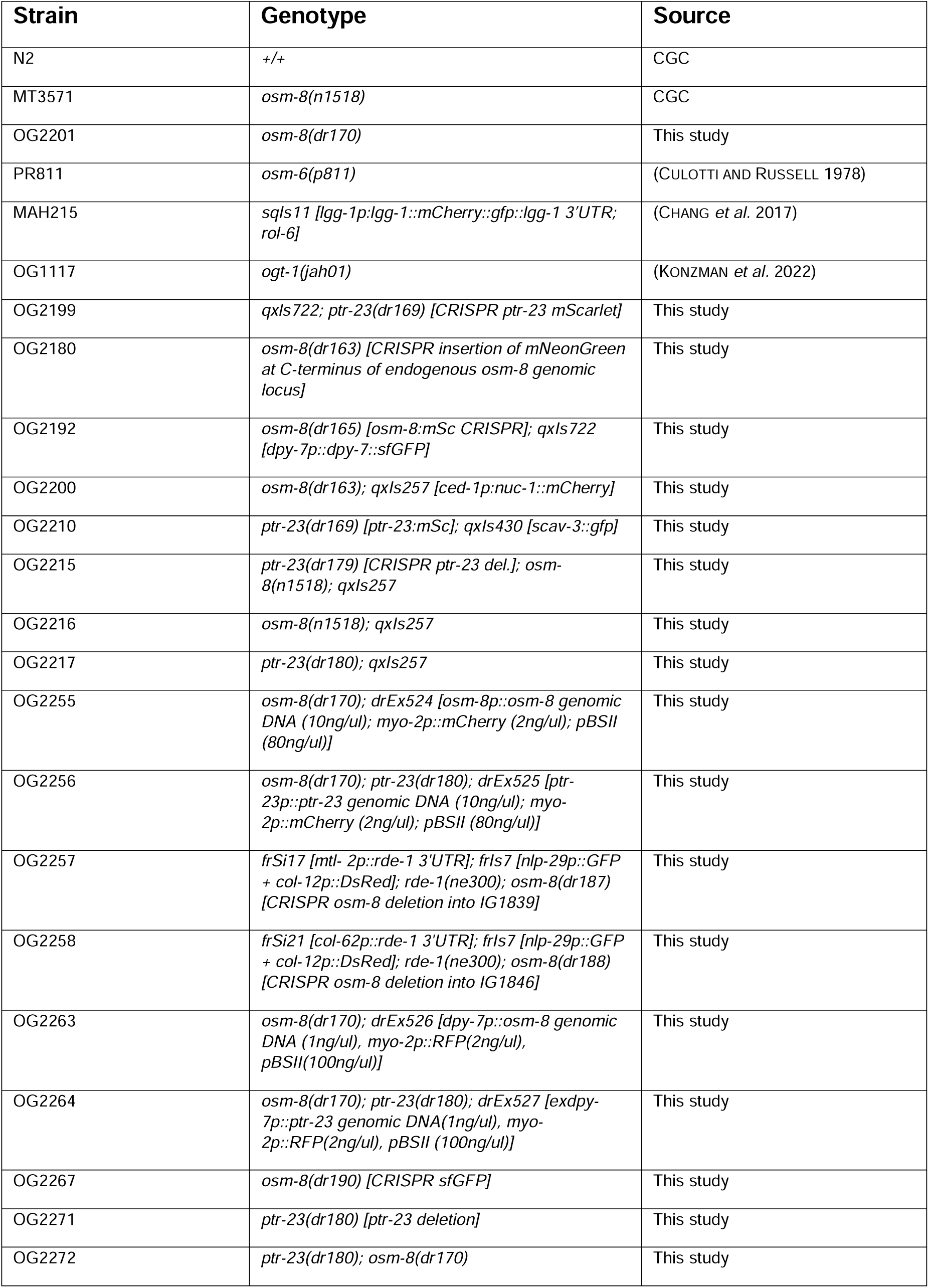
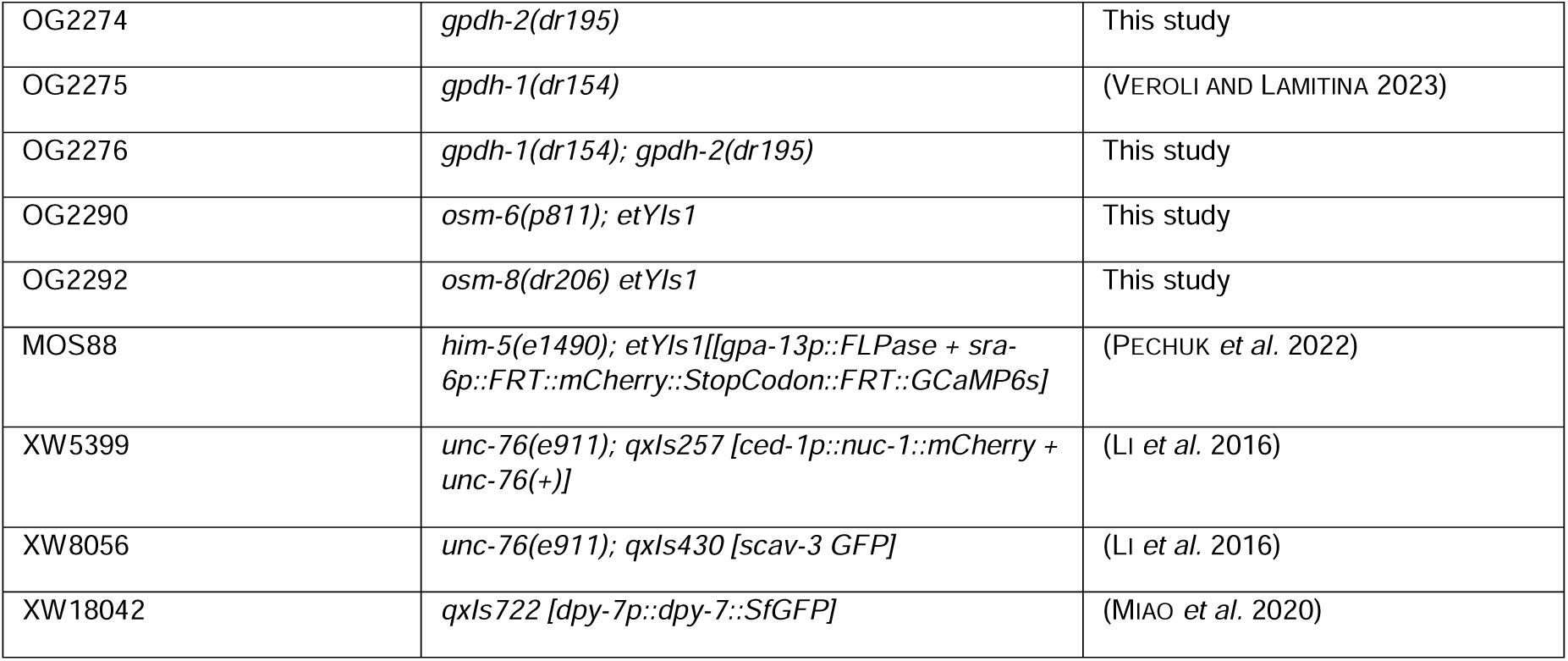
Strains used in this study.

While *osm-8* is annotated to exhibit Osmotic Avoidance (Osm) behavioral defects (www.wormbase.org), these phenotypes have never been described or published. When *C. elegans* encounter hypertonic environments, they first attempt to avoid the stimuli through a well characterized behavioral avoidance response (Culotti and Russell 1978; Bargmann *et al*. 1990; Kaplan and Horvitz 1993; Troemel *et al*. 1995). This avoidance response circuit is primarily driven through detection of osmotic stress via the polymodal sensory neuron ASH. In addition to hypertonicity, ASH also directs avoidance of the volatile odorant 1-octanol via mechanisms that are distinct from those involved in osmosensing (Hart *et al*. 1999). *osm-8* mutants exhibited a strong Osm phenotype similar to that of other Osm mutants, such as *osm-6* (Fig. 1C). Like the Osr phenotype, the Osm phenotype of *osm-8*, was also suppressed by *ptr-23(dr180)* or *ptr-23(RNAi)* (Fig. 1C,D). *ptr-23(RNAi)* did not suppress the *osm-6* Osm phenotype (Fig. 1D), showing that *ptr-23* is a specific suppressor of *osm-8* physiological and behavioral phenotypes.

Many Osm mutants inhibit sensory cilia formation in chemosensory neurons, which leads to characteristic morphological, functional, and behavioral phenotypes. To test if *osm-8* also alters sensory cilia formation, we first investigated the morphology or function of the ASH osmosensory neuron. Unlike *osm-6*, *osm-8* ASH neurons were able to endocytose the extracellular dye DiI, indicating the presence of functional cilia in *osm-8* (Fig. 2A). Next, we determined if the ASH neuron was generally dysfunctional or specifically dysfunctional towards hypertonicity by examining behavioral responses to another noxious chemical detected by ASH – 1-octanol. *osm-8* mutants exhibit a significantly shorter octanol response latency compared to *osm-6,* indicating that the general ability of ASH to respond to stimuli and the downstream neurons mediating avoidance behavior is not strongly affected in *osm-8* (Fig. 2B). Consistent with this, we also found that *osm-8* animals exhibit normal reversal response rates upon direct stimulation of ASH by the light activated channel channelrhodopsin (Fig. 2C). Finally, we compared the behaviors of *osm-6* and *osm-8* in the Osm ring assay (Video S1-S3; (Culotti and Russell 1978)). *osm-6* exhibits relatively few ring encounters because they tend to cross the ring very early in the assay (Fig. 2D,E). In contrast, *osm-8* mutants have significantly more ring encounters and do not cross the ring until much later in the assay as compared to *osm-6* (Fig. 2D,E). Together, these data show that *osm-8* is required for osmotic avoidance behavior and that this phenotype is not due to defects in ASH neuron development or gross abnormalities in the ASH avoidance circuit. However, the kinetics of avoidance in *osm-8* was significantly delayed compared to *osm-6*, suggesting they regulate avoidance through different mechanisms.

**Figure 2.**
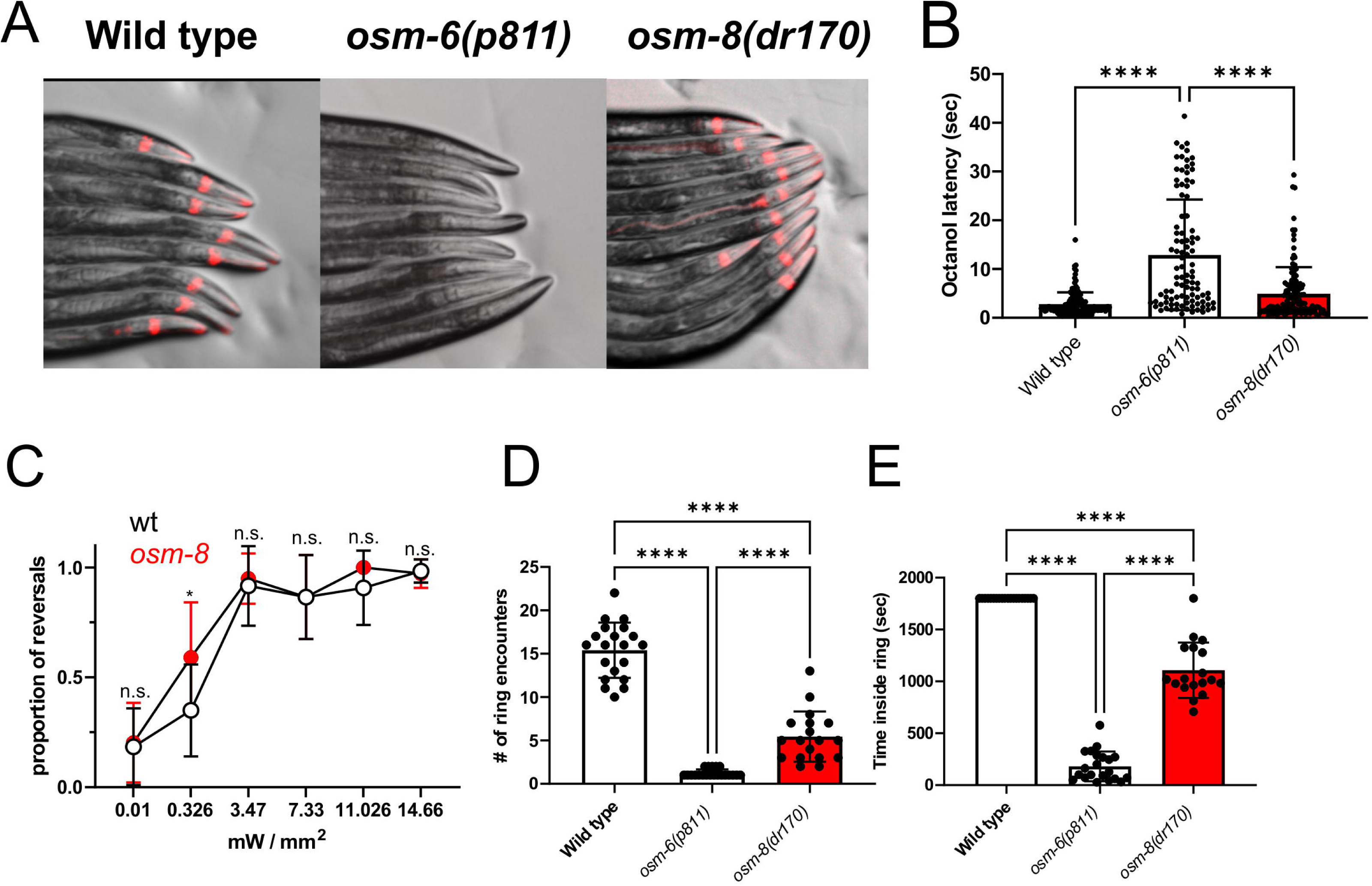
*osm-8* mutants have normal ASH osmosensory neuron development and exhibit a distinct behavioral phenotype from the ciliary mutant *osm-6*. A) DiI staining of day 1 adults of the indicated genotypes. B) Latency to reversal response with 10% octanol. N= 98-136 individual trials per genotype. ****-p<0.0001, One-way ANOVA with Tukey’s multiple comparison test. Individual data points are shown along with the mean ± S.D. C) Proportion of animals reversing in response to ChR2 stimulation in the presence of all-trans retinal. Data shown are the mean +/- S.D. of 9-12 replicates of 5-10 animals each. *-p<0.05, ‘n.s.’ – not significant, One-way ANOVA with Tukey post-hoc testing. D) Number of encounters with the hypertonic ring in the osmotic avoidance assay per individual animal. N=18-21 animals per genotype. ****-p<0.0001, One-way ANOVA with Tukey’s multiple comparison test. Individual data points are shown along with the mean ± S.D. E) Time inside the hypertonic ring in the osmotic avoidance assay per individual animal. N=18-21 animals per genotype. ****-p<0.0001, One-way ANOVA with Tukey’s multiple comparison test. Individual data points are shown along with the mean ± S.D.

The normal responses to 1-octanol and ChR2 stimulation suggest that the behavioral defect in *osm-8* mutants is due to altered ASH osmosensing. If this is the case, ASH neurons should exhibit less neuronal activation in response to hypertonic stimuli than wild type. To test this, we used a microfluidics-based imaging approach and an ASH-specific GCaMP6s reporter to directly measure the calcium responses in the ASH neuron (Chronis *et al*. 2007; Pechuk *et al*. 2022). As a control, we also measured the response of *osm-6* mutants. As previously reported, we found that *osm-6* exhibited no ASH response to a hypertonic stimulus (Philbrook *et al*. 2024)(Fig. 3A-C). In contrast, *osm-8* mutants did respond to hypertonicity, but their response was significantly lower than wild type, although still significantly stronger than *osm-6* mutants (Fig.3A-C). The Osm phenotype of *osm-8* mutants was unaffected by the GCamP transgene (Fig. S1). These data again show that *osm-6* and *osm-8* exhibit substantial phenotypic differences at the level of ASH activation and that the basis of the *osm-8* behavioral defect is likely due in part to the reduced response of ASH to hypertonic stimuli.

**Figure 3.**
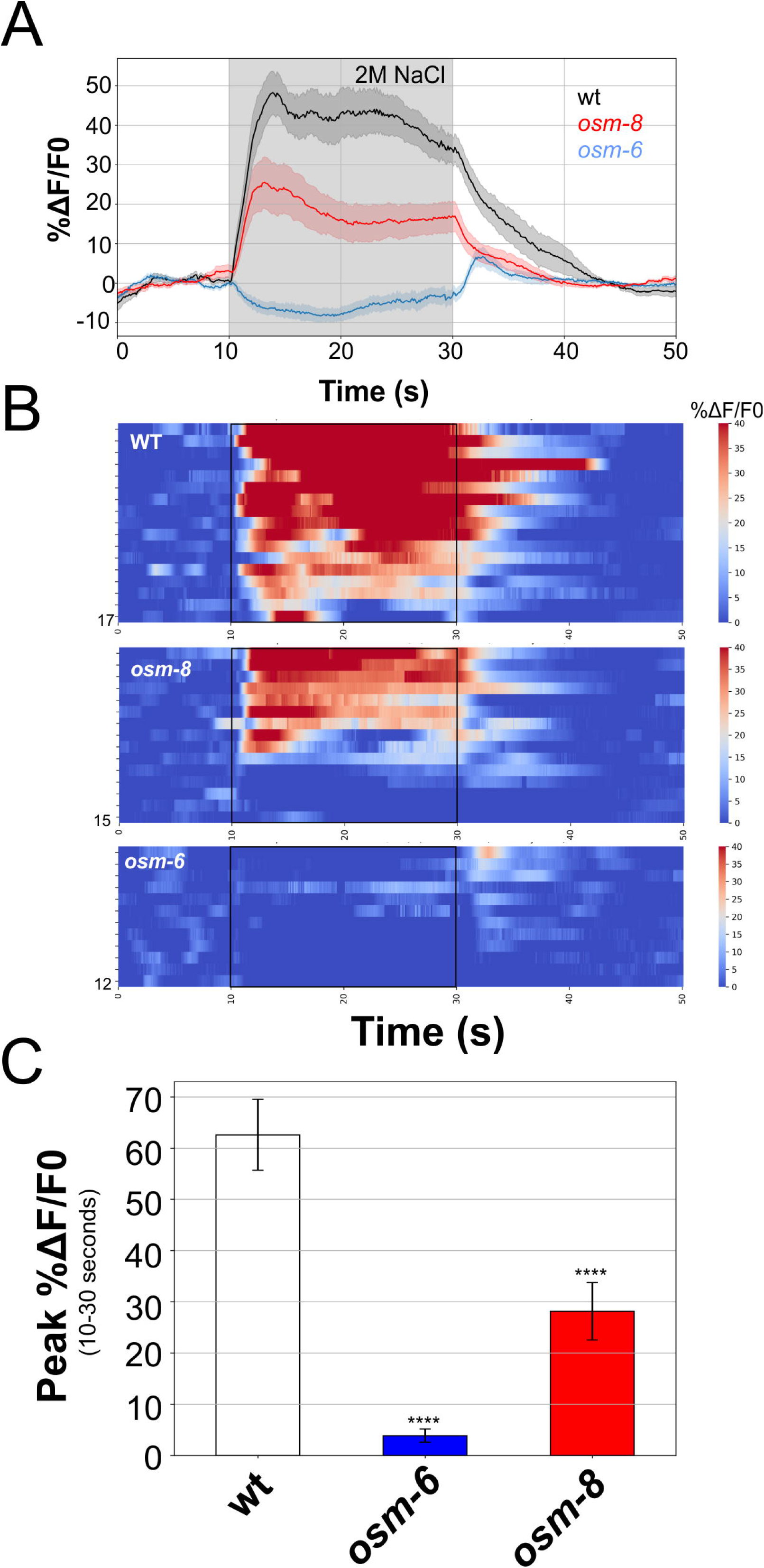
ASH neuronal response to hypertonicity is reduced in *osm-8* mutants. A) Mean +/- SE of wild type, *osm-8*, and *osm-6* GCaMP6s fluorescence in ASH neurons in response to a 20 second exposure to 2M NaCl. N=17, 15, and 12 animals per genotype. Gray shading indicates the period of exposure to 2M NaCl. B) Normalized color coded GCaMP6s calcium responses of ASH to 2M NaCl. Heatmaps show the calcium levels of individual neurons for each genotype. The stimulus is applied from 10-30 seconds. The number of neurons per group is shown at the bottom left corner of each heatmap. C) Mean +/- SEM of the maximum peak %deltaF/F0 during the stimulus period. *** - p<0.001, One-way ANOVA with Tukey post-hoc testing.

There are at least two classes of Osm mutants. Class 1 mutants, such as *osm-6*, are required for the formation of sensory cilia, while Class 2 mutants, such as the TRPV channel *osm-9* (Colbert *et al*. 1997), have normal cilia but are nonetheless functionally defective in the ability to detect all ASH stimuli. Both Class 1 and Class 2 genes are expressed in ASH, as well as other ciliated sensory neurons. When we examined the single cell gene expression map for the *C. elegans* nervous system (Taylor *et al*. 2021), we found that the expression of both Class 1 and Class 2 Osm genes were dramatically enriched in neurons and absent from the hypodermal cells (Fig. 4A). Consistent with previous studies using overexpression reporter transgenes (Rohlfing *et al*. 2011), expression of both *osm-8* and *ptr-23* mRNA exhibit the inverse pattern - they were not detected in sensory neurons, including ASH, but were highly enriched in the hypodermis (Fig. 4A). These expression data suggest the possibility that *osm-8* and *ptr-23* may alter neuronal osmosensing via their function outside of the nervous system.

**Figure 4.**
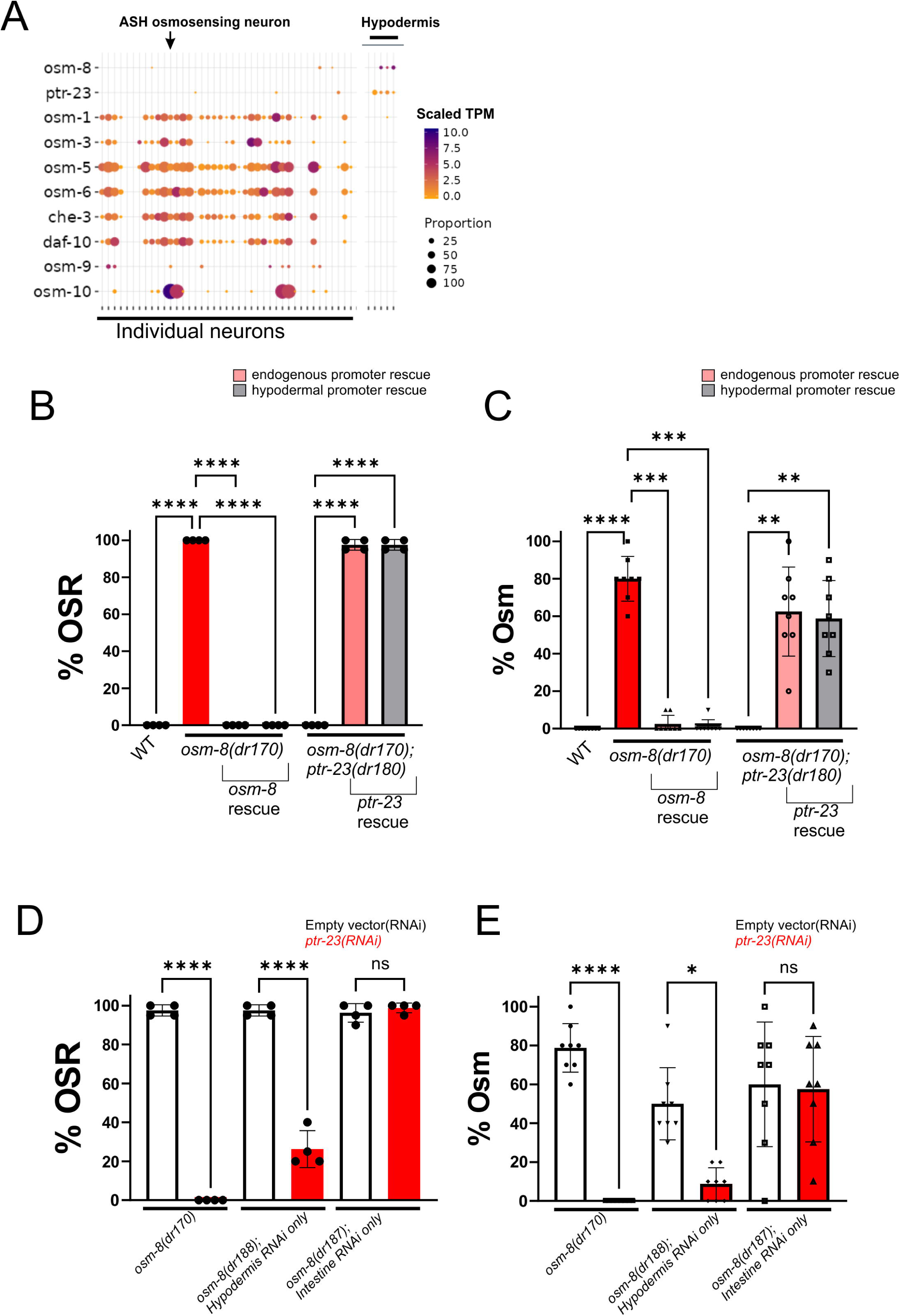
*osm-8* and *ptr-23* hypodermal expression is necessary and sufficient to regulate osmotic avoidance behavior. A) Predicted expression of *osm-8*, *ptr-23*, and other Class 1 & 2 Osm mutants across sensory neurons and hypodermal cells adapted from the *C. elegans* neuronal gene expression map and network (CeNGEN) (Taylor *et al*. 2021) B) Osmotic stress resistance (OSR) phenotype of the indicated genotypes with *osm-8* or *ptr-23* rescue under either their endogenous promoter or the hypodermal specific promoter *dpy-7p*. N=4 replicates per genotype (20 animals per replicate, N=80 per genotype). ***-p<0.001, One-way ANOVA with Kruskal-Wallis post hoc test. Individual data points are shown along with the mean ± S.D. C) Osmotic avoidance phenotype of the indicated genotypes with *osm-8* or *ptr-23* rescue under either their endogenous promoter or the hypodermal specific promoter *dpy-7p*. N=8 replicates per genotype (10 animals per replicate, N=80 per genotype). ****-p<0.0001, *** – p<0.001, **-p<0.01, One-way ANOVA with Kruskal-Wallis post hoc test. Individual data points are shown along with the mean ± S.D. D) Osmotic stress resistance (OSR) phenotype of the indicated genotypes with empty vector or *ptr-23* RNAi. N=4 replicates per genotype (20 animals per replicate, N=80 per genotype). ***-p<0.001, **-p<0.01, One-way ANOVA with Kruskal-Wallis post hoc test. Individual data points are shown along with the mean ± S.D.. E) Osmotic avoidance phenotype of the indicated genotypes with empty vector or *ptr-23* RNAi. N=8 replicates per genotype (10 animals per replicate, N=80 per genotype). ****-p<0.0001, *-p<0.05, ‘ns’-p>0.05, One-way ANOVA with Kruskal-Wallis post hoc test. Individual data points are shown along with the mean ± S.D.

### Hypodermal function of *osm-8* and *ptr-23* is necessary and sufficient for both physiological stress resistance and neuronal stress avoidance behavior

To test if *osm-8* and *ptr-23* act outside of the nervous system to control behavior, we took two approaches. First, we utilized tissue-specific rescue constructs to restore *osm-8* or *ptr-23* expression solely in the hypodermis and examined their ability to rescue *osm-8* or *osm-8;ptr-23* Osm behavioral phenotypes. In *osm-8* mutants, rescue of *osm-8* using either its endogenous promoter or the hypodermal cell specific *dpy-7* promoter fully rescued both the Osr and Osm phenotypes (Fig. 4B,C). Likewise, in *osm-8;ptr-23* mutants, expression of *ptr-23* from either its endogenous promoter or the hypodermal cell specific *dpy-7* promoter restored the Osr and Osm phenotypes (Fig. 4B,C). Therefore, expression of both *ptr-23* and *osm-8* in the hypodermis is sufficient for rescue of neuronal behavioral defects.

Second, we utilized tissue specific RNAi strains in which genes could be inhibited in either the hypodermis or the intestine (Watts *et al*. 2020). Since *osm-8* does not produce a strong or consistent RNAi phenotype (Rohlfing *et al*. 2011), we engineered the *osm-8* deletion into each of these strains using CRISPR/Cas9 and performed *ptr-23(RNAi)*. In a global RNAi background, *ptr-23(RNAi)* fully suppressed the Osm and Osr phenotypes of *osm-8* (Fig. 4D,E). Likewise, when RNAi was restricted to the hypodermis, *ptr-23(RNAi)* also fully suppressed the Osm and Osr phenotypes of the *osm-8* mutant (Fig. 4D,E). However, in an intestine-restricted RNAi strain, *ptr-23(RNAi)* had no effect of any *osm-8* phenotypes (Fig. 4D,E). Taken together, the rescue and tissue specific RNAi data show that *osm-8* and *ptr-23* expression outside of the nervous system in the hypodermis is necessary and sufficient to regulate the physiological Osr and the behavioral Osm phenotypes.

### OSM-8 and PTR-23 are co-localized to hypodermal lysosomes and regulate lysosome abundance and volume

We previously generated a rescuing *osm-8::GFP* transgene and failed to observe any fluorescent signal (Rohlfing *et al*. 2011). Since *osm-8* contains a canonical N-terminal signal sequence, this led us to speculate that OSM-8::GFP might be secreted into a cellular compartment that is recalcitrant to GFP folding and/or fluorescence (Evans *et al*. 2023). To better understand how *osm-8* and *ptr-23* might be regulating behavioral and physiological phenotypes, we generated endogenously tagged alleles using environmentally stable, monomeric fluorophores. Both *osm-8:mNeonGreen::AID*::3XFLAG* and *ptr-23:mScarlett:: AID*::3XFLAG* alleles (referred to as *osm-8::mNG* and *ptr-23:mSc*) were functional, since *osm-8::mNG* did not exhibit an Osr phenotype and *ptr-23:mSc* did not suppress the *osm-8(dr170)* Osr phenotype (Fig. S2). We observed that both OSM-8::mNG and PTR-23::mSc were only expressed in the hypodermis where they exhibited strong co-localization during the L4 stage of development (Fig. 5A-C). However, in adults, OSM-8::mNG was no longer detectable, while PTR-23::mSc was still present (Fig. 5D-F). An identical pattern was observed for both *osm-8:mSc* and *osm-8::sfGFP* CRISPR alleles (Fig. S3). During the L4 stage, PTR-23::mSc co-localized with the lysosomal marker SCAV-3::sfGFP (Fig. 5H-J) while OSM-8::mNG co-localized with the lysosomal marker NUC-1::mCh (Fig. 5K-M). However, PTR-23::mSc was only observed in the lysosomes, suggesting that *osm-8*-dependent signaling occurs through *ptr-23* function in the lysosome.

**Figure 5.**
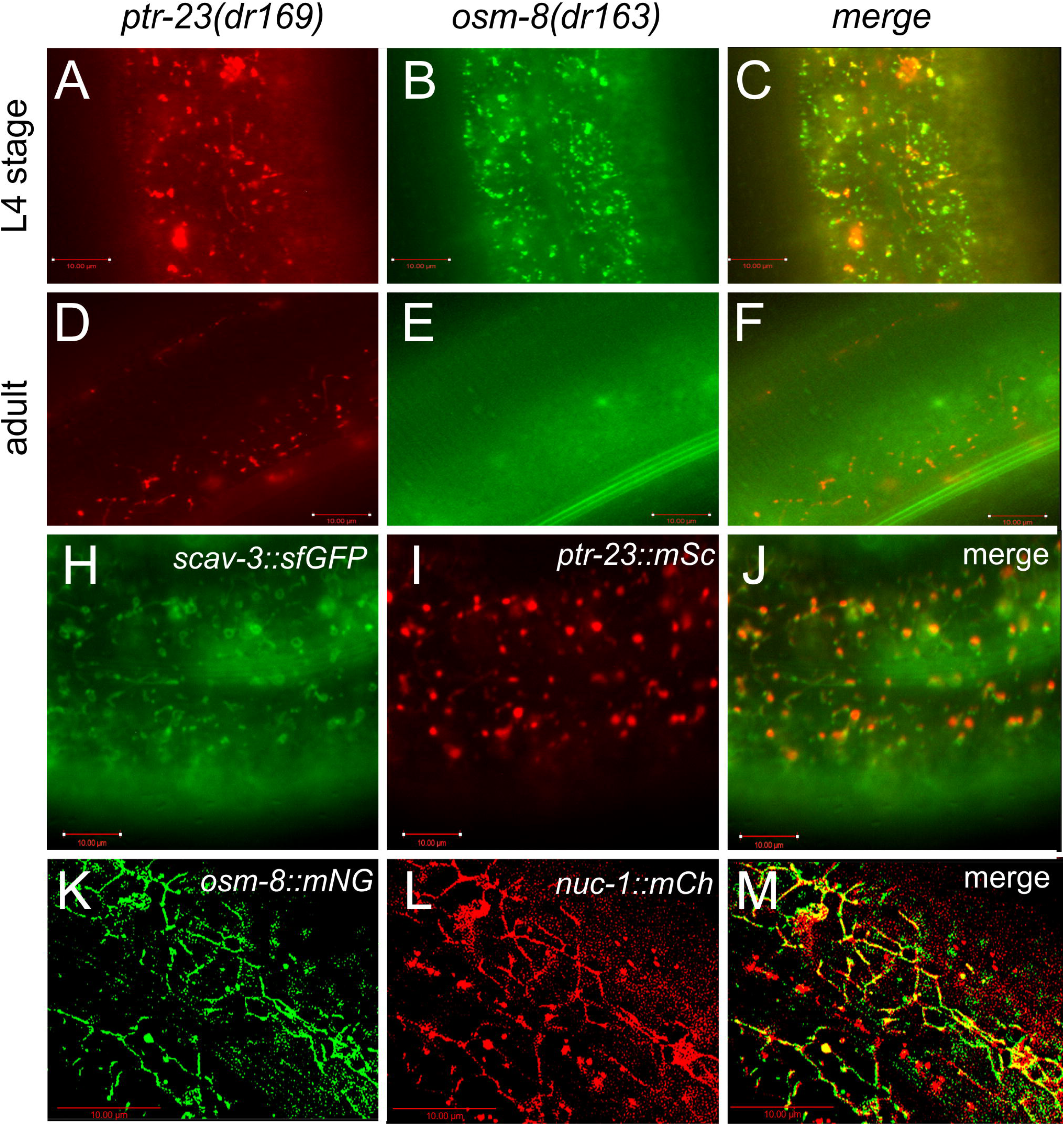
OSM-8 and PTR-23 co-localize in hypodermal lysosomes. A) Wide-field fluorescence images of the hypodermis in L4 stage animals expressing A) *ptr-23::mSc*, *osm-8::mNG*, and C) the merged overlay. Scale bar = 10µm. Wide-field fluorescence images of the hypodermis in day 1 adult animals expressing D) *ptr-23::mSc*, E) *osm-8::mNG*, and F) the merged overlay the indicated endogenously tagged alleles. Scale bar = 10µm. Wide field fluorescence images of Day 1 adults expressing H) the lysosomal marker *scav-3::sfGFP*, I) endogenously tagged *ptr-23:mSc*, and J) the merged overlay. Scale bar=10µm. Deconvolved wide field image of L4 stage animals expressing K) *osm-8::mNG*, L) the lysosomal marker *nuc-1:mCh*, and M) the merged overlay. Scale bar= 10µm. Dotted lines indicate the seam cell boundary, which lack OSM-8 expression.

Given the association of OSM-8 and PTR-23 with the lysosome, which is a major regulator of lipid metabolism (Ebner *et al*. 2025), and the established roles of Patched and Patched-related genes in the transport of hydrophobic molecules including lipids (Zhong *et al*. 2014), we hypothesized that *osm-8* and *ptr-23* may regulate lipid homeostasis. Therefore, we performed unbiased liquid chromatography-high resolution mass spectrometry (LC-HRMS) to identify *osm-8* induced, *ptr-23*-dependent changes in relative lipid abundance. We identified 2,708 distinct lipid species among all 3 genotypes (wild type, *osm-8(dr170)*, *osm-8(dr170); ptr-23(dr180)*). Notably, we found that the lysosome/endosome-specific lipid bis(monoacylglycero)phosphate (also known BMP) was upregulated in *osm-8* mutants but restored to wild type levels in *osm-8; ptr-23* mutants (Fig. 6A,B,C,D). *osm-8* upregulated primarily polyunsaturated BMP species, while monounsaturated species were unaffected (Fig. 6C), indicating some level of specificity in the upregulation of BMP lipids by *osm-8*. Interestingly, the proposed precursor for BMP synthesis, phosphatidylglycerol (PG) (Chen *et al*. 2023; Medoh *et al*. 2023; Bulfon *et al*. 2024; Singh *et al*. 2024), did not increase in *osm-8* but was reduced in *osm-8; ptr-23* (Fig. 6E). Triglycerides, which would be predicted to be elevated in *osm-8* mutants due to the extreme levels of glycerol accumulation, were actually reduced in *osm-8* and this was partially reversed in *osm-8; ptr-23* (Fig. 6F). Similar to previous studies, elevation of BMP by *osm-8* also led to increased cholesterol storage, which is known to mainly occur in the lysosome (McCauliff *et al*. 2019). Increased *osm-8* cholesterol levels were partially reversed in *osm-8; ptr-23* animals (Fig. 6G). Together, these lipidomic data show that *osm-8* and *ptr-23* play key roles in the homeostasis of the lysosomal lipid BMP and accumulation of cholesterol.

**Figure 6.**
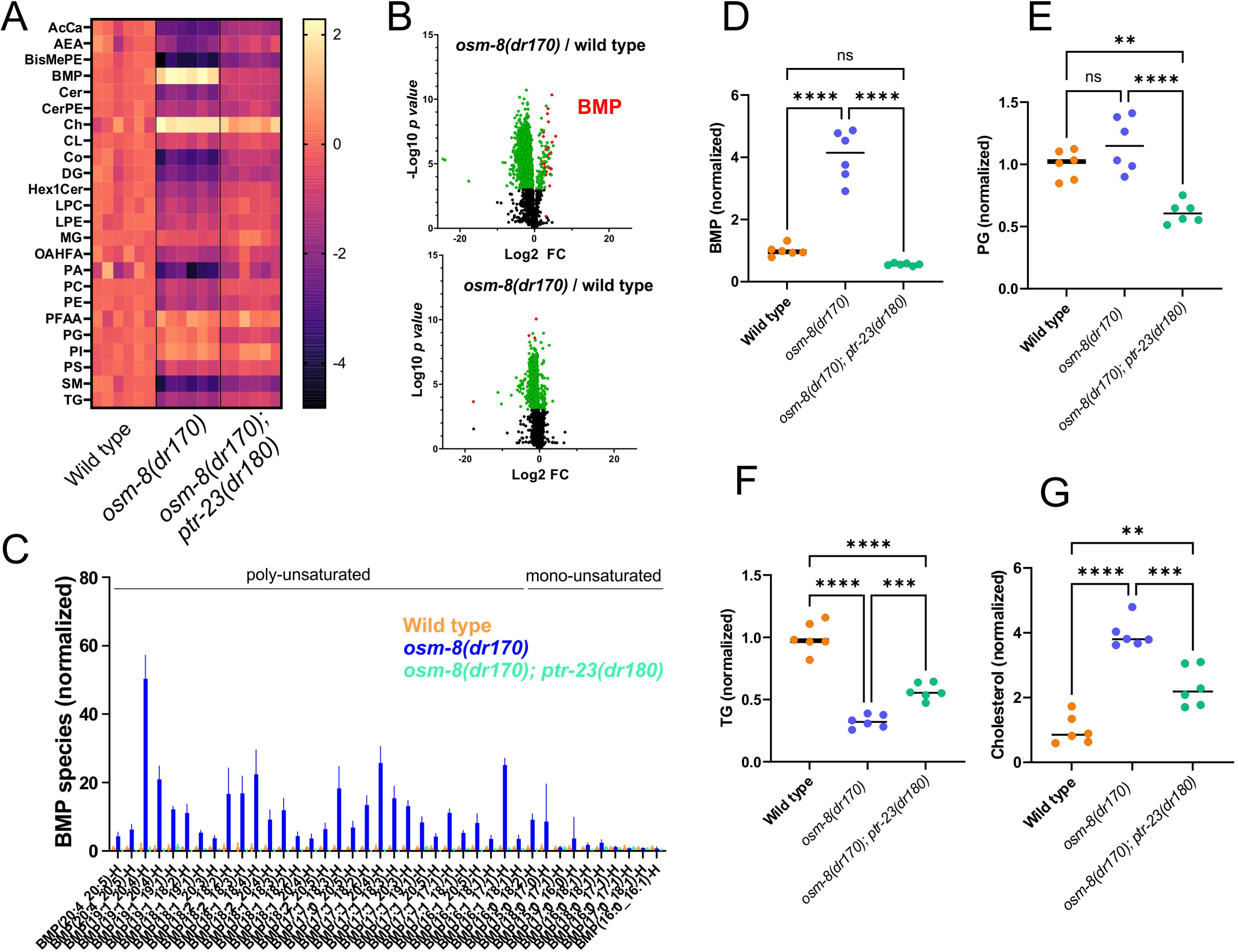
The lysosome-specific lipid bis(monoacylglyerol)phosphate (BMP) and cholesterol are upregulated in *osm-8* mutants in a *ptr-23* dependent manner. A) Heatmap of the normalized log_2_ fold change for each indicated lipid class. Each column represents an individual replicate of 5,000 worms. AcCa=Acyl Carnitine, AEA=N-Acylethanolamine, BisMePE=Bis-methyl phosphatidylethanolamine, BMP=Bis(monoacylglyerol)phosphate, Cer=Ceramide, CerPE=Ceramide PE, Ch=Cholesterol, CL=Cardiolipins, Co=CoenzymeQ, DG=Diglycerides, Hex1Cer=Hexosylceramide, LPC=LysoPC, LPE=LysoPE, MG=Monoglycerides, OAHFA=OAcyl-(gamma-hydroxy)FA, PA=Glycerophosphatidic acid, PC=Glycerophosphocholines, PE=Glycerophosphoethanolamines, PFAA=Primary fatty acid amines, PG=Glycerophosphoglycerols, PI=Glycerophosphoinositols, PS=Glycerophosphoserines, SM=Sphingomyelins, TG=Triacylglycerols. B) Volcano plot of the log_2_ fold change versus log_10_ p-value for all identified lipid species in *osm-8(dr170)* compared to wild type (top) or *osm-8(dr170); ptr-23(dr180)* compared to wild type (bottom). BMP lipids, which are strongly upregulated in *osm-8(dr170)*, are highlighted in red. C) Normalized fold change for all BMP lipid species. Data shown are mean ± S.D. Lipids are organized as poly-unsaturated (>1 acyl chain double bonds) or mono-unsaturated (1 or 0 acyl chain double bonds). D) Normalized log_2_ fold change for BMP lipid group in wild type, *osm-8(dr170)* and *osm-8(dr170); ptr-23(dr180)*. N=6 samples per genotype. Horizontal line indicates the mean. ****-p<0.0001, One-way ANOVA with Tukey post hoc test. E) Normalized log_2_ fold change for phosphoglycerol lipid group in wild type, *osm-8(dr170)* and *osm-8(dr170); ptr-23(dr180)*. N=6 samples per genotype. Horizontal line indicates the mean. ****-p<0.0001, **-p<0.01, One-way ANOVA with Tukey post hoc test. F) Normalized log_2_ fold change for triglyceride lipid group in wild type, *osm-8(dr170)* and *osm-8(dr170); ptr-23(dr180)*. N=6 samples per genotype. Horizontal line indicates the mean. ****-p<0.0001, ***-p<0.001, One-way ANOVA with Tukey post hoc test. G) Normalized log_2_ fold change for phosphoglycerol lipid group in wild type, *osm-8(dr170)* and *osm-8(dr170); ptr-23(dr180)*. N=6 samples per genotype. Horizontal line indicates the mean. ****-p<0.0001, ***-p<0.001, **-p<0.01, One-way ANOVA with Tukey post hoc test.

The elevation in BMP lipids and cholesterol levels observed in *osm-8* suggests that lysosome homeostasis may also be altered in *osm-8*. To test this, we examined lysosomal abundance in the hypodermis of *osm-8*, *ptr-23*, and *osm-8; ptr-23* mutants using the lysosomal marker *nuc-1:mCherry*. We observed a significant increase in the number of hypodermal lysosomes in *osm-8* (Fig. 7A). Automated image analysis revealed that the increase in hypodermal lysosomes seen in *osm-8* was completely eliminated in *osm-8; ptr-23* animals (Fig. 7B). *ptr-23* alone had no effect on the size of the lysosome pool. The increase in the number of lysosomes did not seem to be due to increased autophagy as the abundance of the autophagosomal marker *lgg-1:gfp:mCherry* was not altered by *osm-8* (Fig. 7C). We conclude that *osm-8* and *ptr-23* regulate lysosomal abundance through an autophagy-independent mechanism.

**Figure 7.**
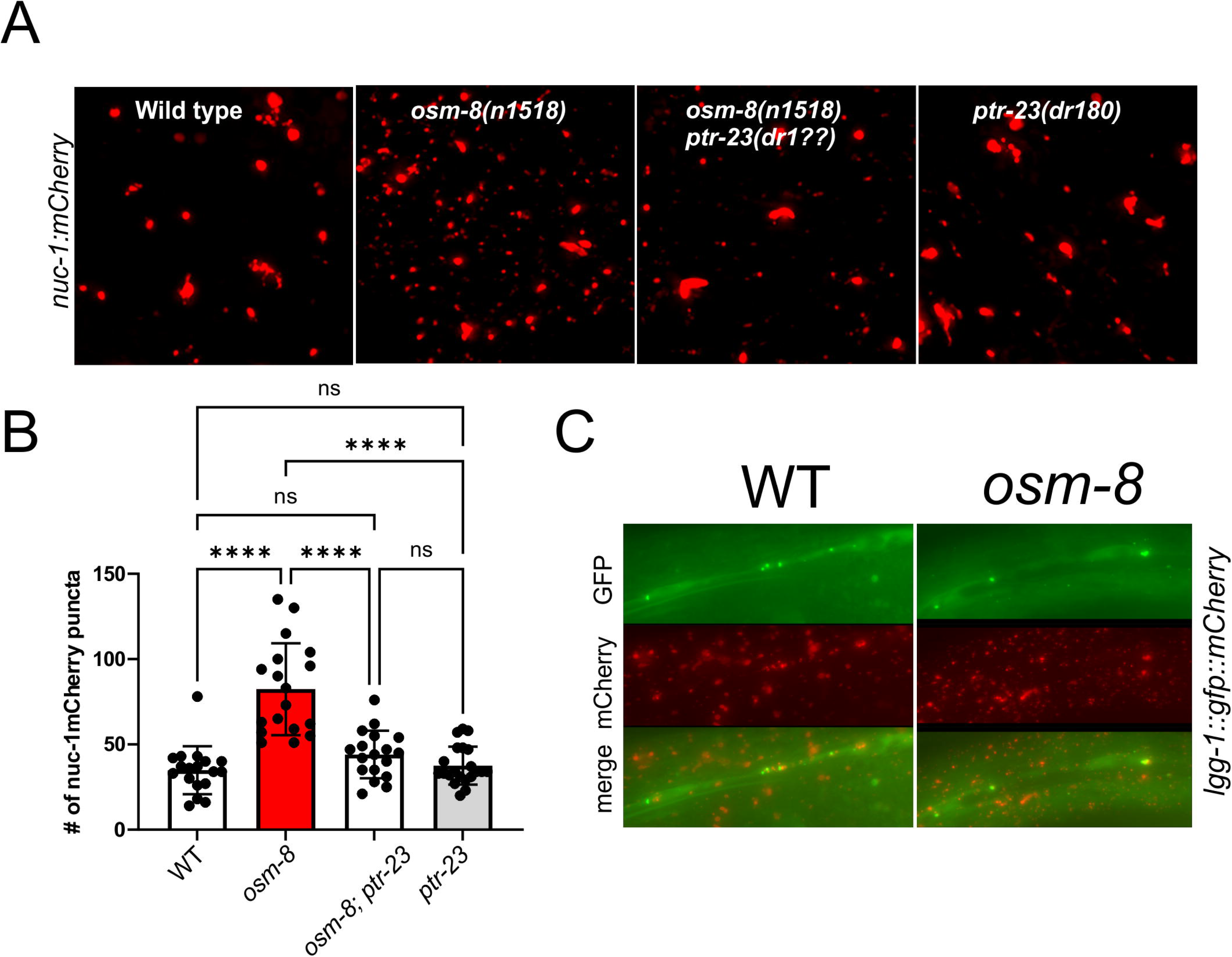
*osm-8* expands the pool of hypodermal lysosomes through a *ptr-23*-dependent, autophagy-independent mechanism. A) Deconvolved flattened Z-stacks of hypodermal *nuc-1:mCh* lysosomal fluorescence in day 1 adult animals of the indicated genotype. Scale bar = 5µm. B) Quantification of the number of *nuc-1:mCh* puncta in each genotype. Individual data points are shown along with the mean ± S.D.. N=18-21 measurements from 6-7 individual animals per genotype. ****-p<0.0001, ‘ns’-not significant, One-way ANOVA with Tukey post hoc multiple comparison testing. C) Widefield images of *lgg-1::gfp:mCh* autophagy marker in day 1 adults of the indicated genotype. Individual green puncta represent autophagosomes while red puncta represent autolysosomes. Scale bar = 5µm

### Acute physiological adaptation of wild type animals to mild hypertonic stress induces a reversible Osm phenotype that requires glycerol biosynthesis and the lysosomal ATPase subunit *vha-5*

*osm-8* mutants genetically activate the same gene expression program seen in wild type animals exposed to non-lethal adaptive levels of hypertonicity (Rohlfing *et al*. 2010) and the expression of many, but not all, of these genes depends on *ptr-23* (Rohlfing *et al*. 2011). We hypothesized that the osmotic avoidance behavioral phenotypes observed in *osm-8* might be replicated in wild type animals by acutely exposing them to mild hypertonic stress. To test this prediction, we exposed wild type animals to 250 mM NaCl for 1, 3, 6, or 24 hours and then performed Osm behavioral ring assays. While non-adapted animals are not Osm, we observed that wild type animals started exhibiting Osm behavior after only 3 hours of adaptation and this increased by 24 hours (Fig. 8A). The acquired Osm behavior in wild type animals was reversed by shifting animals back to 51 mM NaCl and was back to baseline levels after 6 hours (Fig. 8A). Acquired Osm behavior was dose-dependent and could be induced by exposing animal to 150mM NaCl and above (Fig. S4). These data show that the Osm behavior is a physiologically plastic phenotypic trait that is constitutively activated in *osm-8* mutants.

**Figure 8.**
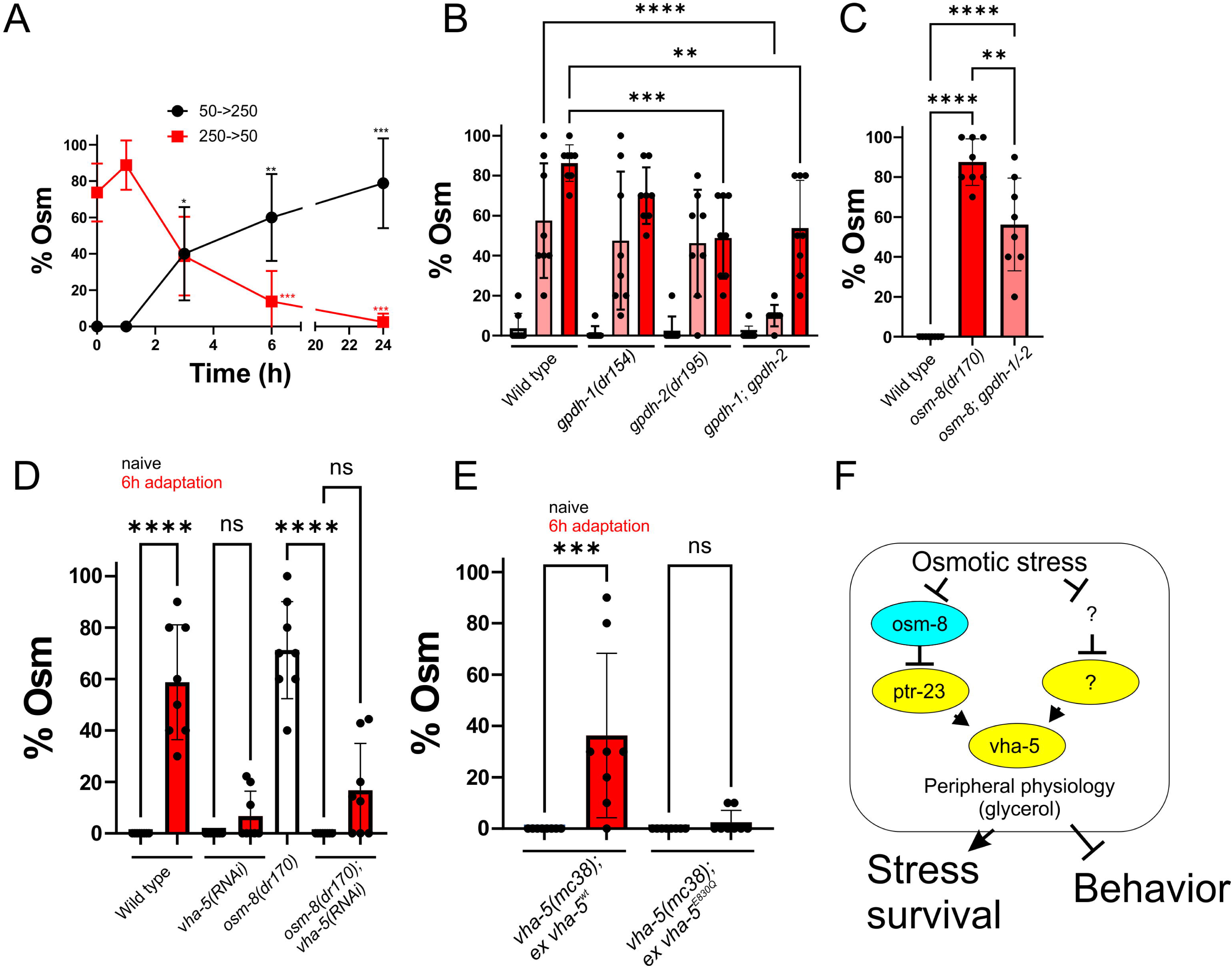
Mild physiological adaptation to osmotic stress drives a reversible Osm phenotype in wild type animals that is dependent on the V-ATPase subunit *vha-5*. A) Adaptation-induced osmotic avoidance behavior in day 1 wild type animals. Black symbols represent animals that were grown on 50mM NaCl and then shifted to 250mM NaCl for the indicated amount of time prior to performance of a standard Osm assay. Red symbols indicate animals that were grown on 250mM NaCl and then shifted back to 50mM NaCl for the indicated amount of time prior to the performance of a standard Osm assay. N=8 replicates per time point for adaptation and reversal (10 animals per replicate, N=80 per genotype). Data points are the mean ± S.D. ****-p<0.0001. ***-p<0.001, ‘ns’-not significant versus t=0. Two-way ANOVA with Tukey post hoc test. B) Osmotic adaptation osmotic avoidance assay in wild type, *gpdh-1(dr154)*, *gpdh-2(dr195)*, and *gpdh-1(dr154); gpdh-2(dr195)* day 1 adult animals. N=8 replicates per genotype and per time point (10 animals per replicate, N=80 per genotype for each time point). **-p<0.01, ***-p<0.001, ****-p<0.0001, One-way ANOVA with Tukey post hoc test. Individual data points are shown along with the mean ± S.D. C) Osmotic avoidance assay in wild type, *osm-8(dr170)*, and *osm-8(dr170); gpdh-1(dr154); gpdh-2(dr195)* day 1 adult animals. N=8 replicates per genotype and per time point (10 animals per replicate, N=80 per genotype for each time point). ‘ns’-not significant, One-way ANOVA with Tukey post hoc test. Individual data points are shown along with the mean ± S.D. D) Osmotic adaptation osmotic avoidance assay in wild type, *vha-5(RNAi), osm-8(dr170),* and *osm-8(dr170); vha-5(RNAi)* day 1 adult animals. N=8 replicates per genotype and per time point (10 animals per replicate, N=80 per genotype for each time point). ****-p<0.0001, ‘ns’-not significant, One-way ANOVA with Tukey post hoc test. Individual data points are shown along with the mean ± S.D. E) Osmotic adaptation assay in vha-5(mc38) day 1 adult null mutant with either *vha-5(+)* wild type rescue or *vha-5^E830Q^*rescue. N=8 replicates per genotype and per time point (10 animals per replicate, N=80 per genotype for each time point). ****-p<0.0001, ‘ns’-not significant, One-way ANOVA with Tukey post hoc test. Individual data points are shown along with the mean ± S.D. F) working model for the role of *osm-8*, *ptr-23*, and *vha-5* in the regulation of peripheral physiology and behavior.

One major physiological consequence of osmotic adaptation in *C. elegans* is accumulation of the organic osmolyte glycerol (Lamitina *et al*. 2004). Genetic deletion of the glycerol-3-phosphate dehydrogenase enzymes *gpdh-1* and *gpdh-2* reduces the accumulation of glycerol by ∼50% (Lamitina *et al*. 2006). To test if glycerol is required for adaptation induced Osm behavior, we adapted *gpdh-1*, *gpdh-2*, and *gpdh-1; gpdh-2* knockouts and analyzed their Osm behavior. We found that Osm behavior was significantly reduced, but not eliminated, in the *gpdh-1; gpdh-2* double mutants, possibly due to the incomplete suppression of glycerol production in this background (Figure 8B). Likewise, the Osm phenotype of *osm-8* was significantly reduced in a *gpdh-1; gpdh-2* mutant background (Fig. 8C). This suggests that both adaptation induced and *osm-8*-induced Osm behavior depends in part on the production of glycerol.

We hypothesized that adaptation induced Osm behavior may require *ptr-23*, since it is required for *osm-8*-induced Osm behavior. However, acquired Osm behavior was not affected by loss of *ptr-23* (Fig. S5). Therefore, we re-examined our prior list of *osm-8* suppressors (Rohlfing *et al*. 2011) and tested if any were required for adaptation induced Osm behavior. We found that the V0 ‘a’ subunit of the V-ATPase *vha-5* was required for both *osm-8* and wild type acquired Osm behavior (6 hour adaptation; Fig. 8D). After 24 hours of exposure to 250mM NaCl, but not 51mM NaCl, all *vha-5(RNAi)* animals died, showing that *vha-5* is also required for hypertonic stress survival (Fig. S6). *vha-5* is only expressed in the hypodermis and excretory cell suggesting that, like *ptr-23*, *vha-5* regulates osmotic avoidance behavior from outside the nervous system (Oka *et al*. 2001; Liegeois *et al*. 2006; Liegeois *et al*. 2007).

Previous studies showed that *vha-5* has both V-ATPase-dependent and -independent functions. The E830Q mutation specifically interferes with the V-ATPase-independent functions in the hypodermis but does not affect the V-ATPase-dependent function in the excretory cell (Liegeois *et al*. 2006). We tested whether *vha-5^E830Q^*was required for *osm-8* phenotypes. We found that *vha-5(+)* but not *vha-5^E830Q^* exhibited an Osr and Osm phenotype in the *osm-8* mutant (Fig. S7A,B). Similarly, *vha-5(+)* exhibited an adaptation-induced Osm phenotype but *vha-5^E830Q^*did not (Fig. 8E). Therefore, the hypodermal-specific V-ATPase independent functions of *vha-5* mediate the effects of *osm-8* on osmotic stress resistance and behavior. Based on our data, we conclude that *osm-8* and *ptr-23* comprise one branch of a multipronged osmotic stress response pathway, which converges on V-ATPase-independent regulation of the V0 subunit *vha-5* and that signaling from the hypodermis via *vha-5* is required to suppress neuronal osmotic avoidance behavior and promote organismal osmotic stress survival (Fig. 8F).

## Discussion

### Peripheral stress resistance pathways drive changes in behavior

Our data show that a behavioral avoidance neuronal circuit can be over-ridden by the activation of stress response pathways in peripheral cells. Genetic or physiological activation of osmotic stress response pathways in the barrier hypodermal cells was sufficient to directly inhibit the osmotic sensitivity of the ASH sensory neuron. This effect was specific for the osmotic avoidance modality of the ASH circuit, since octanol response latency was unaffected. Interestingly, the time course of physiological adaptation and reversal closely matches the accumulation kinetics of the organic osmolyte glycerol and the glycerol biosynthetic enzymes *gpdh-1* and *gpdh-2* were required for physiological adaptation. This suggests an elegant feedback model in which the activation of a physiological response in the peripheral skin tissues produces a metabolite(s) that both blunts the ability of a sensory neuron to detect and avoid threatening environments and enhances organismal survival. Such interoceptive ‘body-brain’ axis signaling between the peripheral digestive system and the brain for the regulation of feeding behavior is well documented (Stover *et al*. 2023). However, our studies of the osmotic stress response and *osm-8* signaling show that such interoceptive signaling is more pervasive than previously appreciated and also takes place outside of the gustatory system.

### *osm-8-ptr-23* regulation of lysosomes may play a key role in communicating physiological state to the nervous system

We showed that *osm-8* and *ptr-23* play essential roles in coordinating stress physiology with behavior. Surprisingly, we find that both genes carry out these functions from the hypodermis. In the case of OSM-8, we find that its accumulation during the L4 stage and disappearance in the adult precisely matches the temporal rescue requirements that we previously defined (Rohlfing *et al*. 2011). While OSM-8 and PTR-23 are localized to lysosomes, regulate lysosome abundance, and control the abundance of lysosome-specific lipids such as BMP, the role of the lysosome in osmotic physiology remains enigmatic. A previous genome wide RNAi screen identified several lysosomal genes as being required for hyperosmotic survival, although *vha-5* was not among the genes identified (Choe and Strange 2008). Another recent study shows that mutation of the cuticular collagen *dpy-7*, which causes an Osr phenotype similar to *osm-8*, exhibits upregulation of some lysosomal genes (Fong *et al*. 2024). One possibility is that lysosomes generate signaling molecules which lead to activation of the osmotic stress response and production of secretory factors that attenuate osmotic avoidance behavior. Such a paradigm was recently described for the role of intestinal lysosomes during aging (Savini *et al*. 2022). Another possibility is that the increase in lysosomal abundance and BMP lipids is a downstream consequence of the activation of glycerol accumulation by *ptr-23*-dependent signaling upon loss of *osm-8*. BMP is the major phospholipid constituent of intralumenal membranes of the late endosome/lysosome, where it regulates sorting and degradation of lipids or lipid cargo (Kobayashi *et al*. 1998). BMP is composed of two monoacylglycerols linked by a phosphate. Elevation of glycerol could drive increased flux through the BMP biosynthesis pathway, resulting in the expansion of BMP enriched lysosomal membranes. The synthesis mechanism of BMP is not well understood, although several enzymes were recently suggested to catalyze different steps in its biosynthesis (Chen *et al*. 2023; Medoh *et al*. 2023; Bulfon *et al*. 2024; Singh *et al*. 2024). Whether or not these genes play important roles in the generation of BMP by the *osm-8/ptr-23* pathway or contribute to the osmotic stress response will require further study.

The V-ATPase proton pump plays a key role in lysosomal physiology by generating the acidic environment required for many aspects of lysosomal function. Like *ptr-23*, the V-ATPase subunit *vha-5* is required for the *osm-8* phenotypes. We were unable to find a consistent role for other Vha genes in *osm-8* phenotypes due to the strong lethal phenotype that was observed in most Vha knockdowns except for *vha-5*. This suggests the possibility that the lysosomal functions of *osm-8*, *ptr-23,* and *vha-5* may not be pH-dependent. *vha-5* has well established V-ATPase-independent roles in *C. elegans* (Liegeois *et al*. 2006). Indeed, our data show that these non-canonical functions of *vha-5* are likely critical since both the Osm and Osr phenotypes of *osm-8* could be suppressed by wild type *vha-5* but not the *vha-5(E830Q)* mutant which preserves canonical V-ATPase functions (Liegeois *et al*. 2006). Non-canonical *vha-5* functions are not well understood but *vha-5* is associated with the secretion of proteins from the hypodermis and serves as a marker of specialized apical plasma membrane structures called meisosomes (Liegeois *et al*. 2006; Aggad *et al*. 2023). Whether or not the meisosomal functions of *vha-5* are required cannot yet be determined since mutations that specifically affect this function(s) are not known.

*ptr-23* is part of the expanded Patched Receptor (*ptc*) and Patched-related (*ptr*) gene family which includes >20 genes in *C. elegans* (Zhong *et al*. 2014). *ptr* genes have established roles in molting and apical extracellular matrix development, as well as the formation of unicellular tube structures (Cohen *et al*. 2021). While Patched and Patched-related proteins are thought to be hydrophobic transporters due to their overall similarity with bacterial resistance-nodulation division (RND) transporters, their specific transport substrates are mostly unknown. One exception is the Patched homolog NPC1, which encodes a lysosomal cholesterol transporter associated with inherited metabolic disorder Niemann-Pick C1 Disease (Saha *et al*. 2020). Mutations in NPC1 cause lysosomal accumulation of cholesterol and elevation of the lipid BMP (McCauliff *et al*. 2019). This causes NPC1 patients to develop a progressive form of ataxia in adolescence and adulthood. Interestingly, we find that mutations in *osm-8* mimic the lipid phenotypes of NPC1 mutations by raising BMP and cholesterol levels. Conversely, we find that mutation of the NPC1-related gene *ptr-23* reverses the *osm-8* induced changes in lipid composition by lowering cholesterol and BMP. Thus, even though PTR-23 localizes to the lysosome and has sequence similarity to the NPC1 gene, it is unlikely to function as a cholesterol transporter. Consistent with this, many of the key residues required for cholesterol transport in NPC1 are not conserved in PTR-23 (Saha *et al*. 2020). Further analysis of the Osr/Osm mutant genetic pathway should help to better understand how OSM-8 and PTR-23 contribute to these dramatic lipidomic and phenotypic changes.

### Physiological signals that drive changes in the nervous system from the hypodermal lysosome

The precise molecular mechanism by which hypodermal osmotic stress signaling represses the osmotic behavioral response remains unclear. Our Ca2+ imaging data suggest hypodermal stress signaling directly inhibits the ASH osmosensory neuron response to hypertonicity. This may be due to the accumulation of the osmolyte glycerol in both osm-8 mutants and wild type adapted animals, which might reduce the osmotic gradient experienced by ASH sensory machinery. Indeed, adaptation-induced behavioral changes precisely track the accumulation and secretion kinetics of whole animal glycerol accumulation (Lamitina *et al*. 2004). Consistent with this model, we find that both *osm-8* and wild type adaptation induced behavioral changes are reduced but not eliminated in animals lacking the biosynthetic enzymes *gpdh-1* and *gpdh-2*, likely due to the fact that these mutants reduce but do not eliminate accumulation of glycerol under hypertonic conditions (Lamitina *et al*. 2006). Other mechanisms could also contribute to the observed behavioral plasticity. For example, a previous study demonstrated that lysosomes generate lipid signals in non-neuronal somatic cells that alter nervous system function and behavior (Savini *et al*. 2022). Given the significant changes in lipid content in *osm-8* mutants, it is possible that such lipid signals may also be involved here. Yet another possibility is that the secretion of several hypodermal expressed neuropeptides, which are upregulated by both *osm-8* and osmotic stress (Rohlfing *et al*. 2010), may communicate with neurons to modify their responses to osmotic stimuli. These neuropeptides are similarly upregulated by infection and were previously shown to regulate infection-induced behavior (Sinner *et al*. 2021). Future studies will be aimed at testing each of these models.

Surprisingly, we found that while loss of *ptr-23* is required for the behavioral adaptation induced by loss of *osm-8*, it is not required for physiological behavioral adaptation induced by high salt. Previously, we showed that even in the absence of *ptr-23*, hypertonicity could still upregulate *gpdh-1* (Rohlfing *et al*. 2011). These data suggests that there is a parallel pathway(s) that regulates the peripheral osmotic stress response and can communicate with neurons. The observation that *vha-5(RNAi)* blocks adaptation induced behavioral changes suggests that this pathway may function in parallel to *ptr-23* and that both pathways converge on *vha-5*. Candidates for such a pathway include the related secreted proteins *osm-7* and *osm-11* (Wheeler and Thomas 2006; Komatsu *et al*. 2008), which may act independently of *ptr-23* (Veroli and Lamitina 2023). Unlike *osm-8*, *osm-7* and *osm-11* are also reported to alter octanol response latency (Singh *et al*. 2011). This suggests that *osm-7* and *osm-11* might modify the ASH avoidance circuit through a different mechanism from *osm-8*. Our findings here show that *osm-8* mutants reduce the response of ASH to a hypertonic stimulus. Whether or not *osm-7* and/or *osm-11* also inhibit ASH responses to hypertonicity is not known. Future studies investigating ASH responses in these mutants will be important to test these models.

In conclusion, our data reveals a non-gustatory ‘body-brain’ axis in *C. elegans* that links peripheral physiological state with behavioral plasticity. The sensation and integration of internal physiological state by the nervous system, a process called interoception, is an emerging but poorly understood process (Khalsa *et al*. 2018). By activating a physiological stress response pathway without altering external environmental input, the *osm-8-ptr-23* pathway in *C. elegans* represents a primitive but tractable interoceptive state in which to define the interoceptive signals that arise from the peripheral tissue, the interoreceptors on neurons, and the mechanisms by which these signals modulate the nervous system. Such mechanisms could provide insights into a wide variety of conditions associated with disruptions in the body-brain axis, such as depression (Avery *et al*. 2014), autism (Hatfield *et al*. 2019), post-traumatic stress syndrome (Simmons *et al*. 2009), and chronic pain (Suarez-Roca *et al*. 2019).

## Materials and methods

### *C. elegans* strains and culture

All strains were cultured at 20°C. A table of all strains used in these studies can be found in Table 1. Wormbase was utilized as the source for all genomic sequence data (Sternberg *et al*. 2024).

### Osmotic stress resistance, adaptation, and osmotic avoidance assays

All assays were performed with Day 1 adults (24 hours post L4 stage). For Osr assays, four replicates of 20 animals were placed in a glass depression slide containing 200µl of M9 with 550 mM NaCl. The number of animals exhibiting movement after 10 minutes was counted.

Osm behavioral assay were performed as previously described (Culotti and Russell 1978), with slight modifications. We printed a ring of 4M NaCl, 0.1% light green SF yellowish dye, pH 7.0 onto standard 3cm NGM plates using a 12mm Falcon snap cap as a template. The cap was gently placed on the agar surface and 40µl of 4M NaCl solution was evenly pipeted around the edge of the cap. The cap was then removed and the 4M NaCl solution was allowed to soak into the plate for 5 minutes prior to the start of the assay. For each assay, 10 day 1 adult animals were placed in S-buffer for ∼1 minute to remove bacteria and then placed in the center of the ring using an eyelash pick. Eight replicates of 10 animals were performed for each genotype and/or condition. After 30 minutes, each plate was scored for the number of animals that crossed the ring. Animals that paralyzed on the ring were censored. To prevent animals from re-entering the ring after crossing, we placed a 10ul spot of OP50 or *empty vector(RNAi)* HT115 bacteria on the outer edge of the plate to which worms navigated and dwelled once they crossed the ring. The presence of food did not result in wild type animals crossing the ring, even after up to 1 hour.

For the analysis of ring encounters and ring dwell time, we collected video (2 frames per second) of the whole 3cm plate over the course of the 30 minute Osm assay using the Wormlab imaging system (N=5-6 animals per video, 4-6 videos per genotype, 18-21 animals total per genotype. Animals were picked into 5µl of S-buffer in the center of the ring to prevent engagement with the ring prior to starting the video recording. After each plate was recorded, worms were tracked using WormLab software throughout the course of the 30 minutes. In order to differentiate an encounter from a crossing, two concentric circular labels were created in WormLab, where one label denoted the inner boundary of the ring and the other label denoted the outer boundary. The sizes of these labels were saved and applied to each video. A worm crossing the first label was denoted as an ‘encounter’, and the same worm crossing the second label was denoted as an escape. The number of encounters a worm underwent was counted, and the time before which the worm escaped the outer label was recorded in seconds. Once a worm crossed the ring, it was no longer considered throughout the remainder of the video. If a worm did not cross the ring after 30 minutes, the time in the ring was denoted as 1800 seconds. Animals that paralyzed inside or on the ring were censored from the analysis.

### Octanol response assay

The 1-octanol avoidance assays were performed as previously published with some modifications (Troemel *et al*. 1995). The morning before conducting the assay, 4µl of OP50 were seeded as 10 separate spots on five 60mm NGM plates per strain and allowed to dry for ∼30 minutes at room temperature. Ten day 1 animals were placed individually onto each spot and left to acclimate to the new plate conditions for approximately 30 minutes. During that time, 1-octanol was diluted to 10% in 100% ethanol. 50µl of diluted 1-octanol was aliquoted into separate PCR tubes for the assays. The 1-octanol aliquots were kept sealed and twelve feet away from the nematodes when assessing 1-octanol avoidance. For the experiments, a fresh horsehair was parafilmed onto the tip of a Q-tip. Then the horsehair was saturated in the diluted 1-octanol, avoiding the parafilm. Then, the horsehair was placed in front of the forward-moving animal. The 1-octanol avoidance response time was recorded upon initiation of backward locomotion. For the subsequent two biological replicates per animal, the horsehair was re-saturated in diluted 1-octanol following the recording of avoidance time to ensure the presence of the odorant throughout the assay.

### Osmotic Survival Assays

Day 1 adults placed onto *empty vector(RNAi)* or *vha-5(RNAi)* plates and incubated for ∼65-72 hr at 20°C. Progeny L4 stage animals were then placed onto either the control or high salt RNAi plates containing 250mM NaCl. 24 hours later, day 1 adults were tested for survival by tapping the plate and by tapping the worm’s body. Animals that moved forward or backward one full body length were scored as alive.

### Imaging and microscopy

L4 stage or day 1 adult animals were immobilized in 50mM levamisole. For images in Figure 4, widefield *nuc-1:mCherry* images were collected as a Z-series (0.27µm step size) on a Leica DMI4000 equipped using a 63x/1.4N.A. lens and a DFC365 digital camera. For quantification of lysosome number, three non-overlapping 25.69µm x 25.69µm x 8.64µm sections centered on the *nuc-1:mCherry* signal in the hypodermis were cropped from each master image. Each cropped section was deconvolved using identical standard settings in imaging software (Leica AF). Puncta in the 3D deconvolved cropped Z-stacks were then analyzed using Aivia software. Training took place on one wildtype image stack and equivalent threshold settings were then applied to all image stacks for all genotypes to generate object counts.

### Microfluidics and Ca^2+^ imaging

.Custom microfluidic devices were based on a previous design with modification (dual layers to facilitate sample loading/removal and a narrower 50µm diameter worm trap tapered to a 15µm opening) (Chronis *et al*. 2007). Silicone wafer molds were printed using maskless photolithography at the University of Pittsburgh Nanofabrication Core Facility. Following silianization of the molds, PDMS elastomer (1:10) was mixed, degassed, poured onto silicon wafers, and incubated at 65°C for 2-4 hours. The polymerized PDMS was removed from the wafer, cut into individual devices, and 1mm access ports were punched. Chips were sealed to cleaned glass coverslips with a 1 minute plasma treatment of the contact surfaces under vacuum and then applying gentle pressure to seal the PDMS to the coverslip. Sealed devices were baked at 80°C for 30 minutes to enhance bonding.

Devices were connected to four fluid reservoirs via a direct PVC tubing connection. Stimulus flow was controlled through a valve control system (Valvebank 4, Automate Scientific). Stimuli were diluted in filtered Milli-Q water. Images were acquired with a Leica DMi8 inverted microscope with a 40x dry objective (0.6NA) and a Leica K5 sCMOS camera at 10Hz with 4×4 binning (100ms exposure time). Day 1 adult animals grown without starvation for at least 2 generations were picked onto a clean NGM plate to remove bacteria, aspirated from the clean plate in 0.3mM levamisole to minimize movement in the device, and loaded into the device trap. Prior to imaging, each animal was allowed to equilibrate in the device with flow for 3 minutes without illumination. Each animal was then subjected to 20 seconds of control, 20 seconds of stimulus, and 20 seconds of control solution (600 frames). Data were collected over at least two days using the same solutions with some animals from each genotype being collected each day. Only a single ASH neuron was recorded from each animal.

For GCaMP analysis, images stacks were first aligned using the ‘Template Matching’ plugin in Fiji. An ROI was manually drawn around the ASH soma and a matching ROI was placed outside the worm for measuring background fluorescence. Mean neuron soma and background fluorescence measurements were used for analysis using custom Python scripts. First, the background subtracted fluorescence was calculated for each neuron. Next, frame numbers were converted to time (ie 10 frames per second). F0 was then calculated for each neuron as the average background subtracted fluorescence during the 15-20 second period. The delta F/F0 was then calculated for each neuron using the formula (F-F0) / F0 where F is the background-subtracted F at each time point. To correct for GCaMP photobleaching, an exponential decay function was fit to the deltaF/F0 data for each neuron during the 10-20 second and 50-60 second periods and this curve was subtracted from the deltaF/F0 data for each neuron. Finally, the corrected deltaF/F0 values for each neuron were converted to percentages and this data was used for time plots, heat maps, and peak value plots. For all time data, the first 10 seconds of data is omitted since this period often contained sample positioning and focusing artifacts. Peak %deltaF/F0 was calculated as the maximum change in fluorescence in the 20-40 second stimulus period. All code is available at: https://github.com/LamitinaLab/Witrado_et_al_2025.git

### Transgene rescue experiments

*osm-8* and *ptr-23* genomic fragments containing the promoter (∼2Kb), coding sequence, and 3’ UTR (∼1Kb) was PCR amplified (Q5 polymerase, New England Biolabs) and cloned into the pMinT2.0 vector. To derive clones with hypodermal-specific expression, we replaced the *osm-8* or *ptr-23* promoters with a *dpy-7* promoter using recombination-based cloning (High-fidelity assembly, NEB). The sequence of all plasmids was verified by whole-plasmid DNA sequencing (Plasmidsaurus).

To generate *osm-8* rescue lines, we microinjected *osm-8(dr170)* with a mix containing pBluescript II (80ng/µl), pCFJ90/*myo-2p::mCherry* (2ng/µl), and the *osm-8* rescue plasmid (20ng/µl or 2ng/µl for the *dpy-7p* plasmid, which was toxic at higher concentrations). To generate *ptr-23* rescue lines, we injected *osm-8(dr170); ptr-23(dr180)* with a mix containing pBluescript II (80ng/µl), pCFJ90/*myo-2p::mCherry* (2ng/µl), and the *ptr-23* rescue plasmid (20ng/µl or 2ng/µl for the *dpy-7*p plasmid, which was toxic at higher concentrations). Stable extrachromosomal array lines that segregated *myo-2p::mCherry* in the progeny were selected for analysis.

### CRISPR/Cas9 genomic editing

The *osm-8* deletion allele *dr170* and the *ptr-23* deletion allele *dr180* were generated using previously described CRISPR/Cas9 editing methods (Veroli and Lamitina 2023). mNeonGreen and mScarlett tags were amplified from pJW2171 and pJW2098 (Ashley *et al*. 2021) and inserted into the endogenous *osm-8* or *ptr-23* genes using our standard CRISPR/Cas9 methods (Urso *et al*. 2023). While these alleles contain an AID* degron tag, we did not observe a reduction in either mNeongreen or mScarlett signals in the presence of 1mM K-NAA and a globally expressed TIR1 (*wrdSi58*), likely due to the AID* degron tag being located in the lysosome lumen, which would be inaccessible to the nucleocytoplasmic TIR1. All alleles were verified by DNA sequencing. A table of all primers used in this study can be found in Table 2.

**Table 2.**
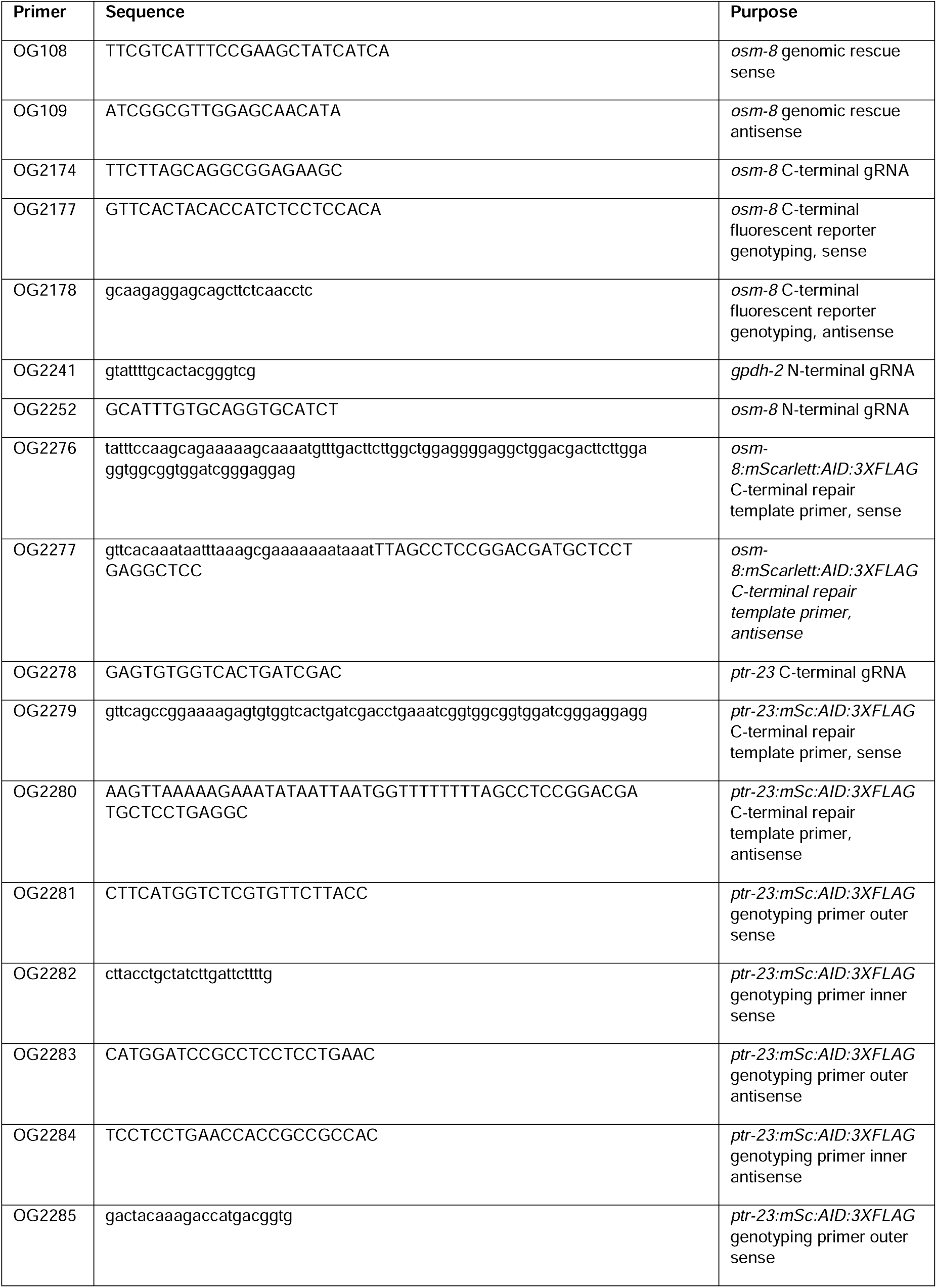
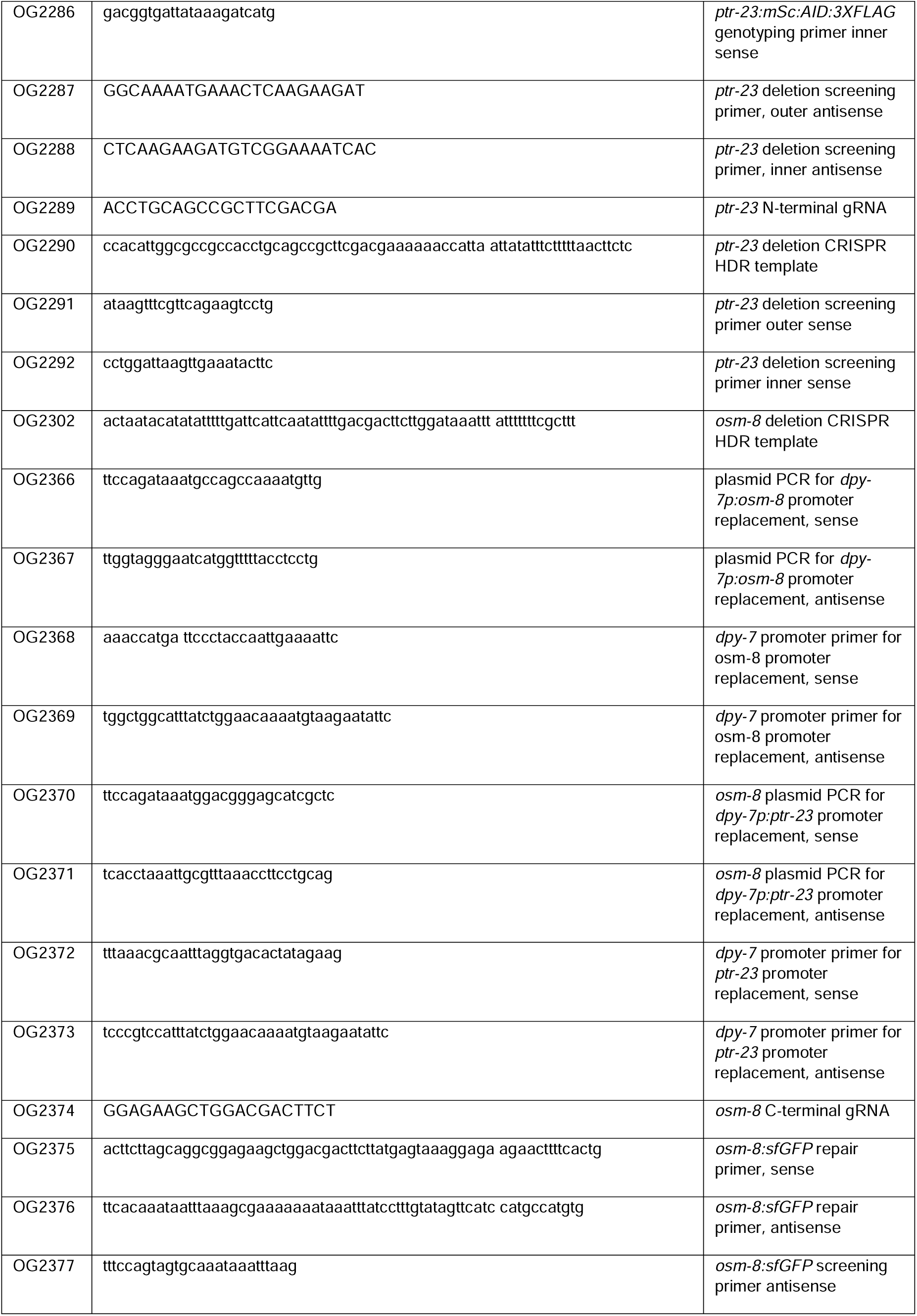
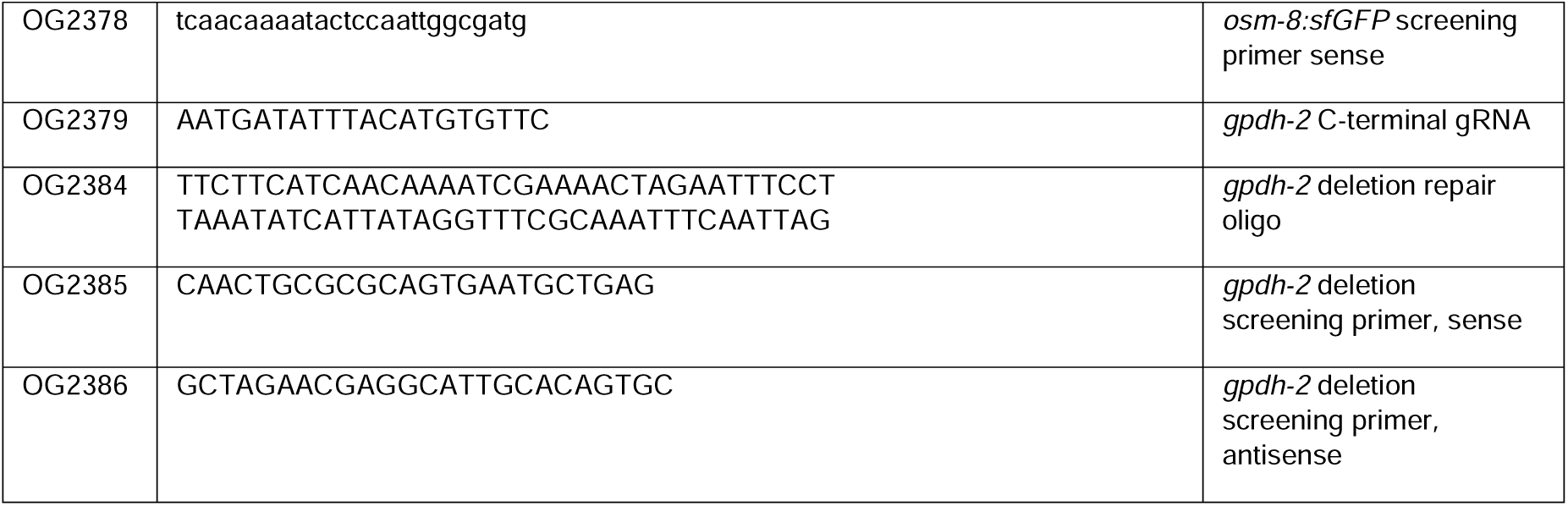
Oligos used in this study.

### Lipidomics

For each replicate, N2, *osm-8(dr170)*, and *osm-8(dr170); ptr-23(dr180)* animals were seeded on 3×15 cm NGM plate with a lawn of OP50 as starved L1s. Animals were grown at 20°C until animals were day 1 adults. We collected a total of six replicates for each genotype for analysis. Worms were washed off plates using M9 buffer, pelleted in a swinging bucket centrifuge at 2,000xg for 1 minute, and then washed three times. Worm pellets were resuspended in 10ml of M9 buffer and incubated at 20°C with rocking for 15 minutes to evacuate OP50 bacteria from the intestine. Worms were pelleted and the density of worms per microliter was calculated for each strain. For each sample, ∼5,000 worms were pipeted into 2.0ml tubes containing a MP Bio Matrix A (garnet/ceramic sphere) and frozen at −80°C prior to processing.

Metabolic quenching, lysis, and lipid extraction was performed by adding 0.5mL ice cold phosphate buffered saline. Samples were homogenized in MP Biomatrix A tubes at 60 Hz for 1 minute. Uncleared supernatant (400 µL) was transferred to a clean glass tube containing 10 µL of LipidSplash, deuterated internal standards (Avanti Polar Lipids. Alabaster, AL) and subjected to Folch extraction. Samples were rested on ice for 10 minutes before phase separation via centrifugation at 2500 x g for 15 minutes. The organic phase (0.7 mL) was transferred to a clean glass vial and dried to completion under N2. Samples were resuspended in 100 µL of 1:1 acetonitrile:isopropanol and 3µL was injected for online LC-HRMS analysis.

Briefly, samples were injected via a Thermo Vanquish UHPLC and separated over a reversed phase Thermo Accucore C-18 column (2.1×100mm, 5μm particle size) maintained at 55°C. For the 30 minute LC gradient, the mobile phase consisted of the following: solvent A (50:50 H2O:ACN 10 mM ammonium acetate/0.1% acetic acid) and solvent B (90:10 IPA:ACN 10 mM ammonium acetate/0.1% acetic acid). The gradient started at 30% B and increased to 43%B over 2 minutes followed by an increase to 55%B in 0.1 minute. The organic increased to 65%B over the next 10 minutes, followed by an increase to 85%B over 6 minutes. For column washing the gradient increased to 100%B over 2 minutes and was held for 5 minutes followed by 5 minutes of equilibration at 30%B. The Thermo ID-X tribrid mass spectrometer was operated in both positive and negative ESI mode. A data-dependent MS2 method was used with scanning in Full MS mode from 200 to 1500 m/z at 120,000 resolution with an AGC target of 5e4 for triggering MS2 fragmentation using stepped HCD collision energies at 20, 40, and 60% in the Orbitrap at 15,000 resolution. Source ionization settings were 3.5kV and 2.4kV spray voltage, respectively for positive and negative mode. Source gas parameters were 35 sheath gas, 5 auxiliary gas at 300°C, and 1 sweep gas. Calibration was performed prior to analysis using the PierceTM FlexMix Ion Calibration Solutions (Thermo Fisher Scientific). Internal standard peak areas were then extracted manually using Quan Browser (Thermo Fisher Xcalibur ver. 2.7), normalized to weight and internal standard peak area, then graphed using GraphPad PRISM (ver 9.0). Untargeted differential comparisons were performed using LipidSearch 4.2 (Thermo Fisher) to generate a ranked list of significant lipid compounds at the class and species-specific levels.

## Statistical Analysis

Comparisons of means were analyzed with either a two-tailed Students t-test (2 groups) or ANOVA (3 or more groups) using either the Dunnett’s, Sidak, or Tukey’s post-test analysis as indicated in GraphPad Prism 9 (GraphPad Software, Inc., La Jolla, CA) or Python 3 libraries. p-values of <0.05 were considered significant.

## Data Availability Statement

Strains and plasmids are available upon request. The authors affirm that all data necessary for confirming the conclusions of the article are present within the article, figures, and tables. All raw data for all figures is available via FigShare.

## Acknowledgements

This work was supported by NIH R01GM135577 to T.L. and NIHS10OD023402 to S.G.. Some mutant strains were provided by the CGC, which was funded by the NIH Office of Research Infrastructure Programs (P40 OD010440). We thank Jun Chen for work performed in the University of Pittsburgh Nanofabrication and Characterization Core Facility (RRID:SCR_05124). NFCF services and instruments used in this project were graciously supported, in part, by the University of Pittsburgh. We also thank Piali Sengupta, Ph.D. and Jamie Yeon, Ph.D. (Brandeis University) for guidance and advice on Ca^2+^ imaging.

## Conflict of Interest Statement

The authors have no conflicts of interest to declare

## Supplemental Figures

Figure S1 Osmotic avoidance behavior in wild type and *osm-8* mutant animals with or without the *etYIs1* GCaMP6s transgene. N=8 replicates per genotype (10 animals per replicate, N=80 per genotype). ****-p<0.0001, ‘ns’-not significant, One-way ANOVA with Tukey post hoc test. Individual data points are shown along with the mean ± S.D.

Figure S2 Osmotic stress resistance phenotype of tagged *osm-8* and *ptr-23* alleles. N=4 replicates per genotype (20 animals per replicate, N=80 per genotype). ****-p<0.0001, ‘n.s.’ – not significant, One-way ANOVA with Tukey posthoc testing.

Figure S3 Localization of endogenously tagged *osm-8* alleles. A) *osm-8(dr165) [mScarlet* CRISPR allele] B) *dpy-7::sfGFP* C) merge. Scale bar = 10 microns. D) *osm-8(dr190) [sfGFP* CRISPR allele] in L4 or E) day 1 adult animals. Scale bar = 10 microns.

Figure S4 Dose response of osmotic adaptation-induced Osm behavior in wild type animals. Adaptation period was 24 hours. N=8 replicates per genotype and per time point (10 animals per replicate, N=80 per genotype for each concentration). ****-p<0.0001, ‘ns’-not significant, One-way ANOVA with Tukey post hoc test. Individual data points are shown along with the mean ± S.D.

Figure S5 Adaptation induced Osm behavior in wild type and *ptr-23(dr180)* mutants. N=8 replicates per genotype and per time point (10 animals per replicate, N=80 per genotype for each time point). ****-p<0.0001, ‘ns’-not significant, One-way ANOVA with Tukey post hoc test. Individual data points are shown along with the mean ± S.D.

Figure S6 Survival of *vha-5(RNAi)* day 1 adults in either wild type or *ptr-23(dr180)* after 24 hour exposure to NGM plates with either 50mM NaCl or 250mM NaCl. N=4 replicates per condition (20-25 animals per replicate, 80-100 animals per condition per genotype). ****-p<0.0001, ‘ns’-not significant, One-way ANOVA with Tukey post hoc testing. Individual data points are shown along with the mean ± S.D.

Figure S7 Non-canonical V-ATPase independent functions of *vha-5* are required for *osm-8* phenotypes. A) OSR phenotype of wild type or *osm-8* mutants in the *vha-5* null mutant *mc38* with rescue from either a *vha-5(+)* wild type or *vha-5E830Q* mutant transgene. N=4 replicates per genotype (20 animals per replicate, N=80 per genotype). ****-p<0.0001, ‘n.s.’ – not significant, One-way ANOVA with Tukey post hoc testing. B) Osm behavior in wild type or *osm-8* mutants in the *vha-5* null mutant *mc38* with rescue from either a *vha-5(+)* wild type or *vha-5E830Q* mutant transgene. N=8 replicates per genotype and per time point (10 animals per replicate, N=80 per genotype for each time point). ****-p<0.0001, ‘ns’-not significant, One-way ANOVA with Tukey post hoc test. Individual data points are shown along with the mean ± S.D.

Video S1. Wild type osmotic avoidance behavior. Ring assay with 4M NaCl. Spot of OP50 *E. coli* in upper right quadrant provides both a motivating stimulus to cross the ring and a ‘sink’ to retain worms that cross the barrier to prevent them from re-entering the ring. Note that wild type animals exhibit chemotaxis towards the OP50 stimulus even when inside the ring, consistent with an odorant-based attractive mechanism.

Video S2. *osm-6(p811)* osmotic avoidance behavior. Ring assay with 4M NaCl. Spot of OP50 *E. coli* in upper right quadrant provides both a motivating stimulus to cross the ring and a ‘sink’ to retain worms that cross the barrier to prevent them from re-entering the ring. Note that *osm-6* mutants do not appear to chemotaxis towards the OP50 stimulus, consistent with their general defect in both soluble and odorant-based sensory neuron function.

Video S3. *osm-8(dr170)* osmotic avoidance behavior. Ring assay with 4M NaCl. Spot of OP50 *E. coli* in upper right quadrant provides both a motivating stimulus to cross the ring and a ‘sink’ to retain worms that cross the barrier to prevent them from re-entering the ring. Note that unlike *osm-6*, *osm-8* mutants chemotaxis towards the OP50 stimulus, consistent with the hypothesis that *osm-8* mutants do not cause a general defect in chemosensory function.

## Notes

https://github.com/LamitinaLab/Witrado_et_al_2025.git

### Competing Interest Statement

The authors have declared no competing interest.

### Summary of Updates

Microfluidics-based calcium imaging of ASH neuron added as new Figure 3. Supplemental Figures S1-7 and Videos S1-3 added. Abstract updated to include new data. Methods section updated. Author list and affiliations updated.

