## Supplementary figures and images for "Soma to neuron communication links stress adaptation to stress avoidance behavior"

### Fig S1

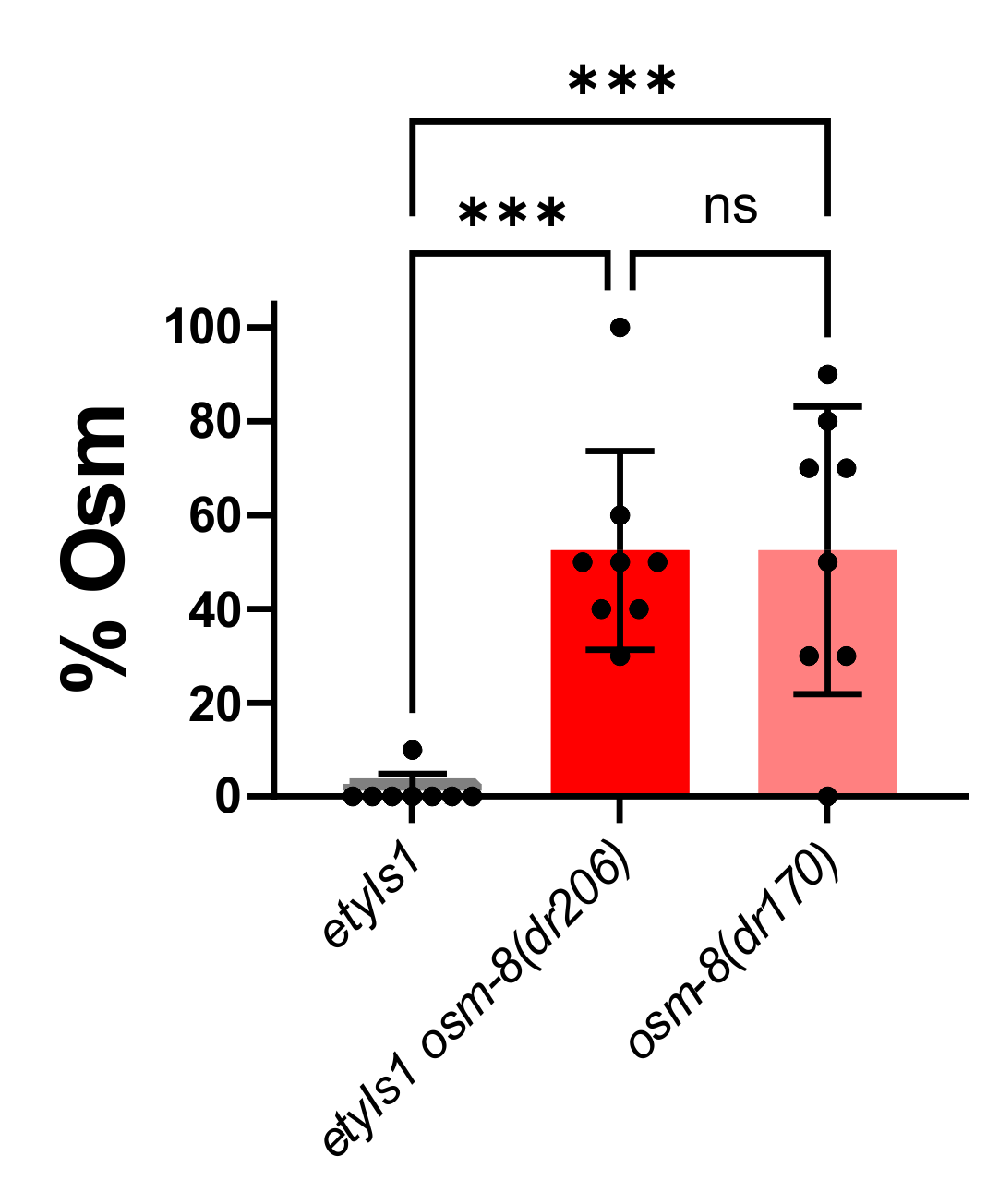

### Fig S2

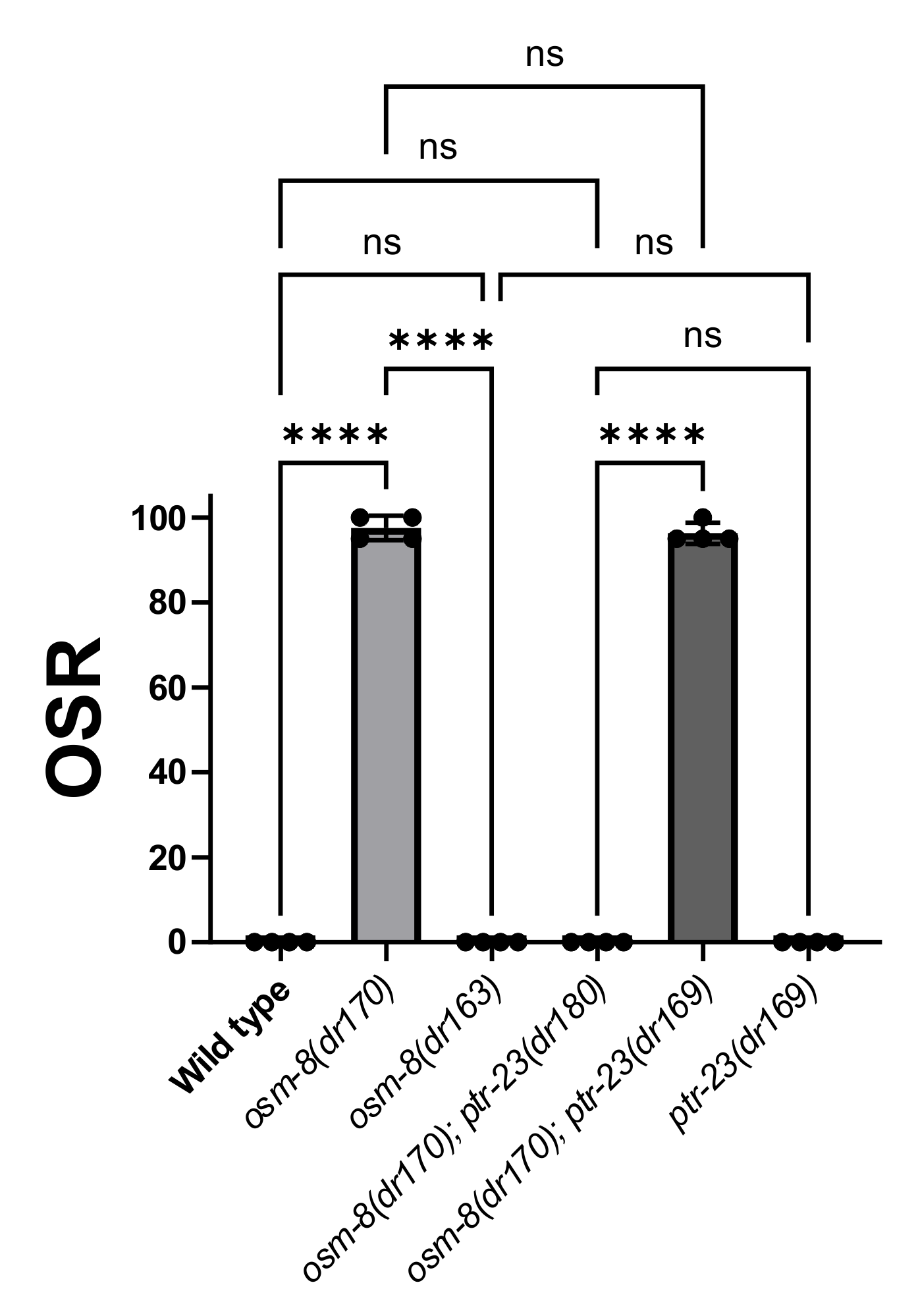

### Fig S3

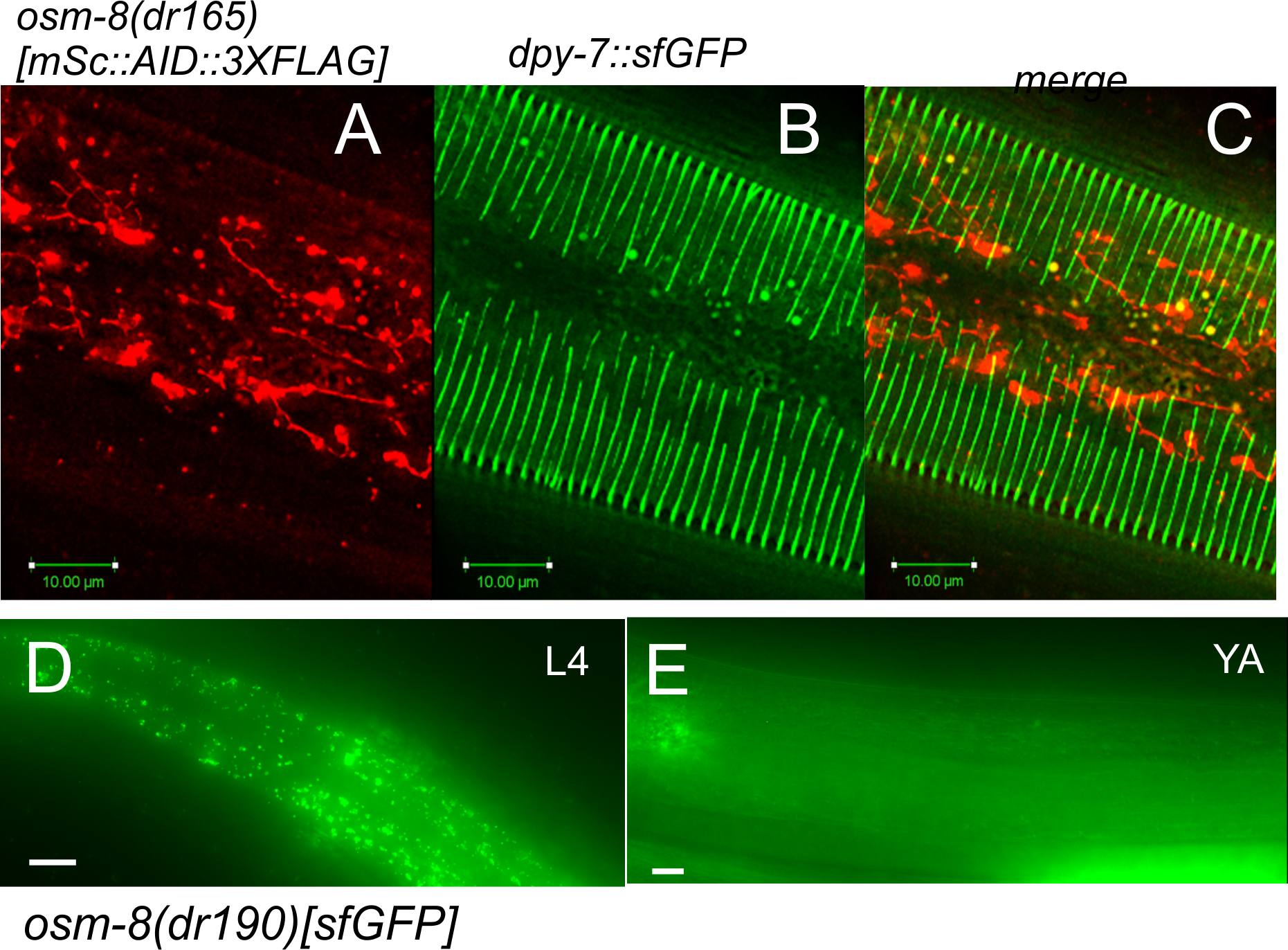

### Fig S4

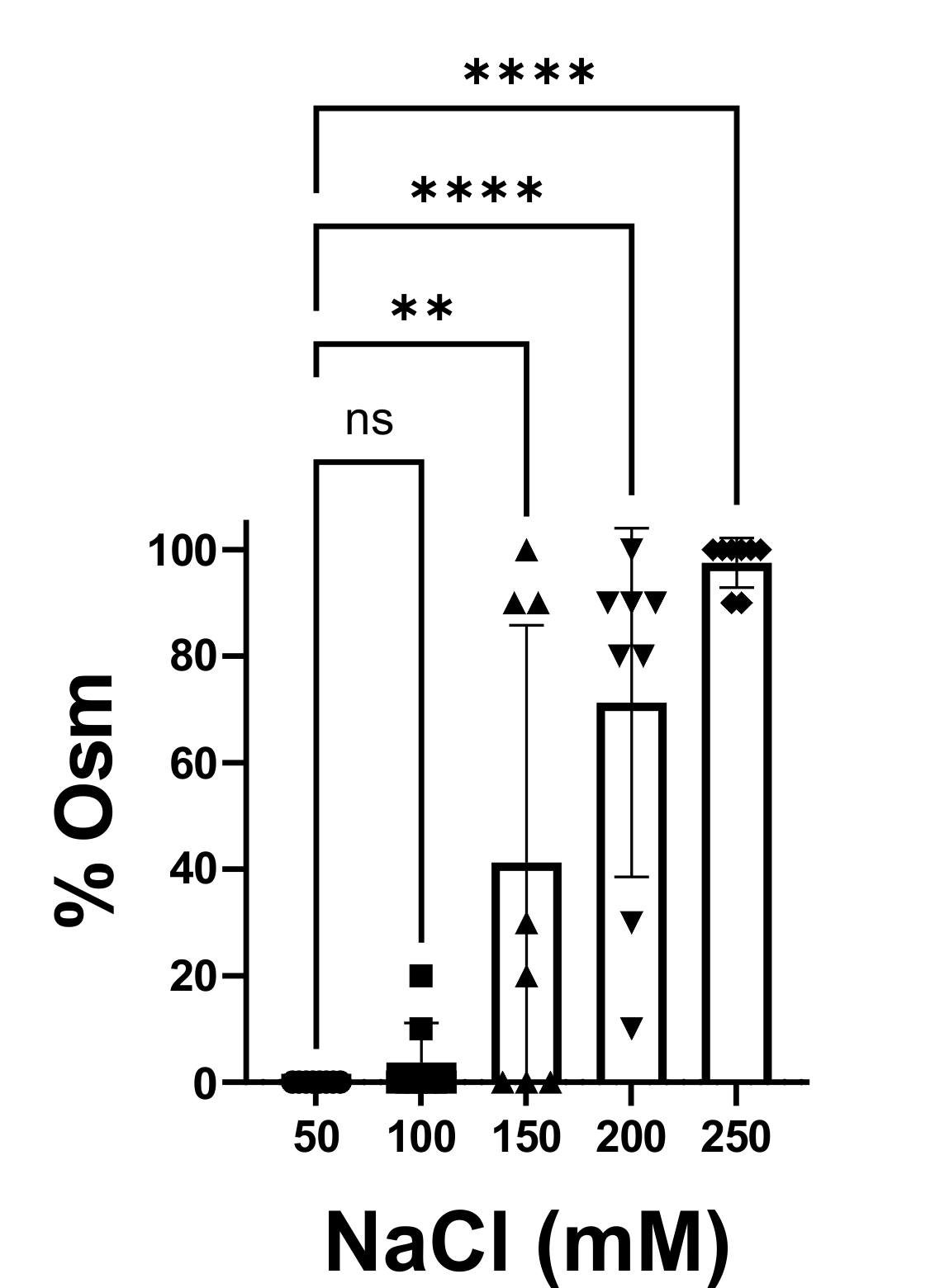

### Fig S5

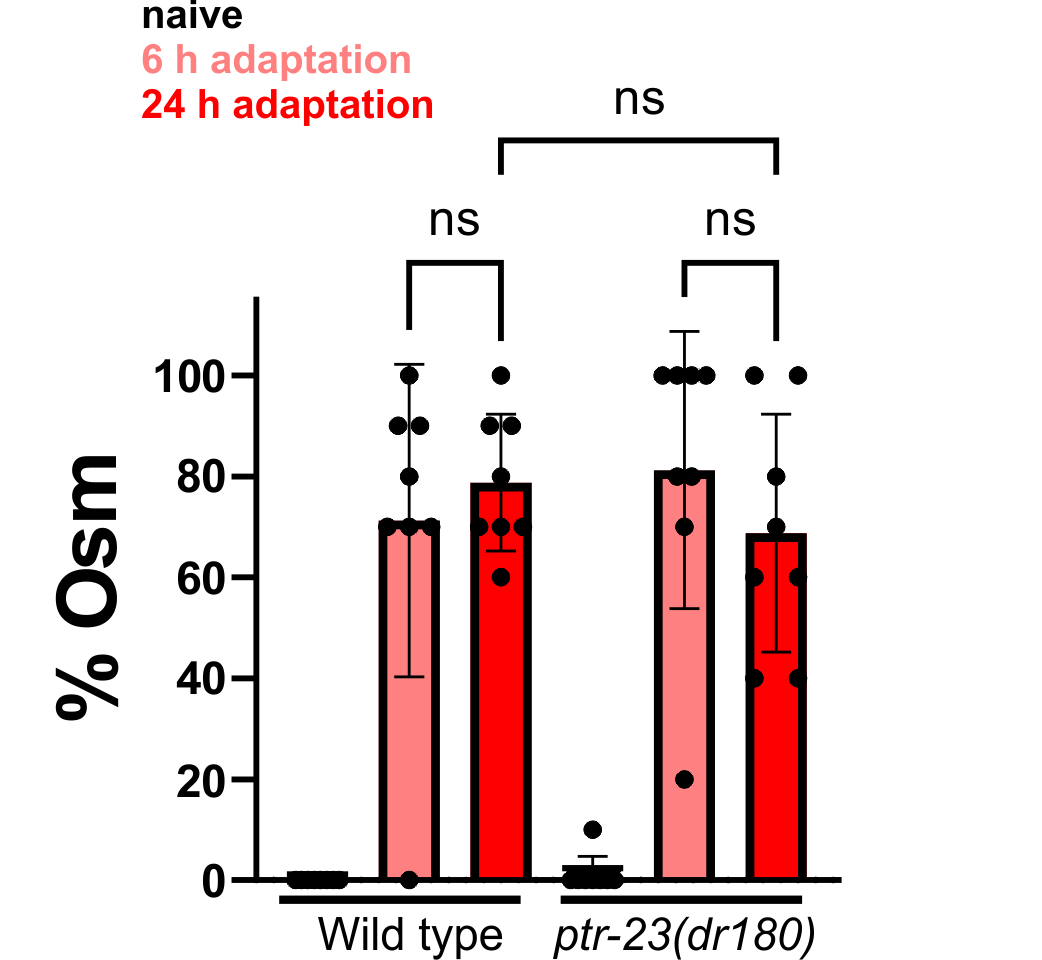

### Fig S6

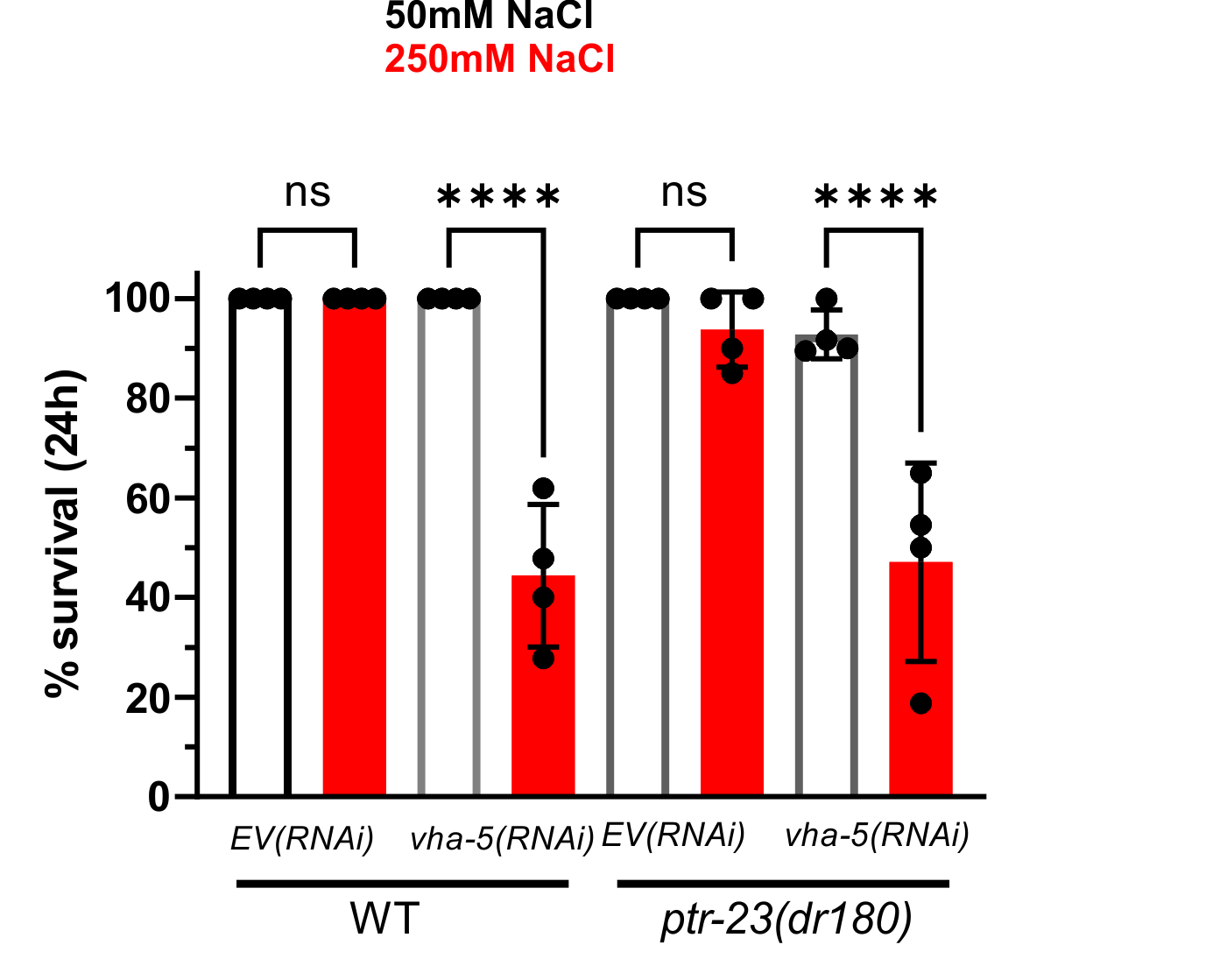

### Fig S7

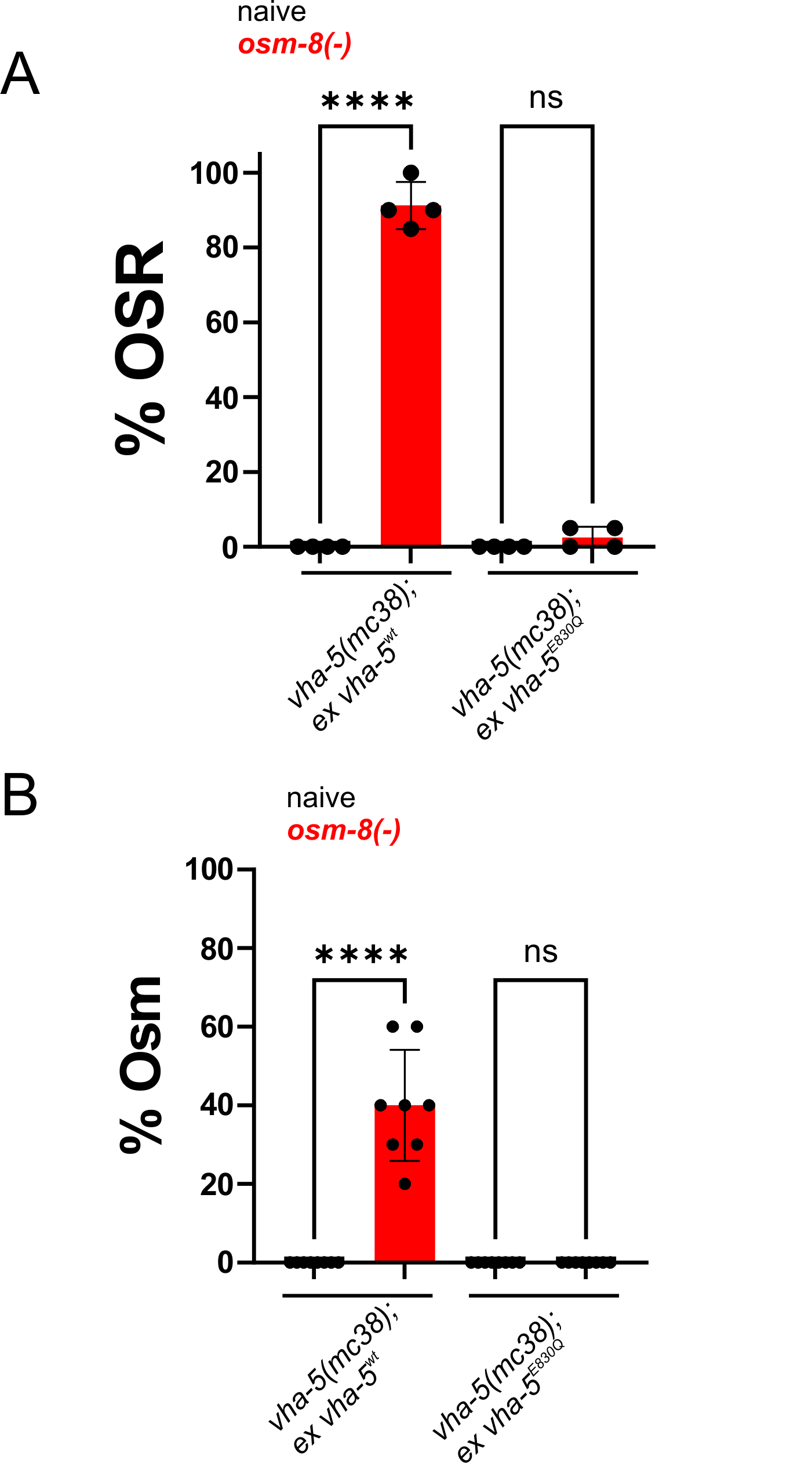
